# On an algorithmic definition for the components of the minimal cell

**DOI:** 10.1101/333682

**Authors:** Octavio Martínez, M. Humberto Reyes-Valdés

## Abstract

Living cells are highly complex systems comprising a multitude of elements that are engaged in the many convoluted processes observed during the cell cycle. However, not all elements and processes are essential for cell survival and reproduction under steady-state environmental conditions. To distinguish between essential from expendable cell components and thus define the ‘minimal cell’ and the corresponding ‘minimal genome’, we postulate that the synthesis of all cell elements can be represented as a finite set of binary operators, and within this framework we show that cell elements that depend on their previous existence to be synthesized are those that are essential for cell survival. An algorithm to distinguish essential cell elements is presented and demonstrated within an interactome. Data and functions implementing the algorithm are given as supporting information. We expect that this algorithmic approach will lead to the determination of the complete interactome of the minimal cell, which could then be experimentally validated. The assumptions behind this hypothesis as well as its consequences for experimental and theoretical biology are discussed.

## Introduction

It is clear that some cell components are essential for survival, while others, at least under certain conditions, are dispensable [1]. Classical examples of the former are non-redundant genes coding for components of the DNA replication machinery [2], while examples of the latter are genes or proteins involved exclusively with secondary metabolism [3]. Classification of cell elements into these separately defined categories has been carried out within all domains of life, ranging from prokaryotes such as *E. coli* [4], to humans [5], and there is a database exclusively devoted to essential genes [6], which current version includes also noncoding genomic elements [7].

Even when the determination of essential cell components has been biased toward genetic elements [8], the recognition of the fact that the concurrent presence of non-genomic elements is indispensable for cell survival resulted in the concept of ‘minimal cell’, which began with the pioneering efforts to construct artificial cells in the 1960s [9], and advanced to form the field of synthetic biology [10]. On the other hand, the determination of the smallest set of components that can sustain life has obvious importance for a solid foundation of biology, and will help in the understanding of critical cellular processes [7, 11, 12].

It is important to underline that the definition of ‘essential cell components’, genomic or otherwise, depends to some extent on particular environmental conditions [13], e.g., in a bacteria with a mutation affecting the synthesis of an amino acid ‘*x*’, such amino acid will be classified as ‘essential’ only when it is absent form the culture media. However, if we take a functional view, it appears impossible to avoid the fact that, for example, an element to synthesize RNA from a DNA template (an RNA polymerase) is essential for all free-living cells.

### Experimental approaches

Experimental approaches to determine minimal gene sets began, before the genomic era, by generating random gene knockouts and determining which of them were lethal [14]. In general, estimation of the number and nature of essential genes can proceed by different methods of genome-wide gene inactivation in both, prokaryotes and eukaryotes. The estimated proportion of essential genes ranges from less than 6% in *C. elegans*, up to almost 80% for *Mycoplasma* species (summarized in [13]). For a human cancer cell line, the authors in [5] infer that approximately 9.2% of the genes are essential. Interestingly, this proportion is relatively close to the estimate for *C. elegans* (6%), and appears to indicate that complex organisms have a lower percentage of essential genes, or in other words, that a larger proportion of their genomes is concerned with tasks not completely essential for cell function. However, those tasks could be indispensable for survival at the organism’s level.

Another possibility to infer the essentiality of genes is provided by comparative genomics. The general argument of this approach is that orthologous genes conserved in genomes separated by very large periods of independent evolution, should be indispensable for cell function; however, this set must be completed by genes that perform an indispensable function, but are non-orthologous (nonorthologous gene displacement; NOD) [15].

A third experimental strategy to determine essentiality is the artificial synthesis of a genome. In this regard, the pioneering experiments by Craig Venter and his team [16], built a bacterial genome *in vitro* and transplanted it into a different (but closely related) species, resulting in what the press called “the world’s first synthetic life form”. In the Venter group’s experiment, after a few generations all proteins in the receptor species were synthesized from the information present in the transplanted genome. The achievements of Venter’s group in transplanting prokaryote genomes–which generated strong public interest and scientific controversy [17], have been followed by the synthesis of a functional eukaryotic chromosome from yeast [18], and then by the design and construction of more than one third–approximately 6.5 of the total 16 chromosomes, with the aim of producing a synthetic genome for this organism (*Saccharomyces cerevisiae*) within the ‘Sc2.0’ project [19–25]. Scientists within this program set up the BioStudio software framework [19] to design the yeast chromosomes, with rules that included the removal of repetitive regions and introns, the substitution of the TAG stop codon by TAA, the relocation of transfer RNA genes into a neochromosome, and the introduction of loxPsym sites at the 3’ ends of nonessential genes to induce genome rearrangements [26]. This last manipulation allows the selection of phenotypes and their corresponding genotypes by the inducible evolution system “SCRaMbLE” [19]. The ‘final wet-lab model’ for a cell, will be an engineered system in which all components are obtained by *in vitro* synthesis and assembly. In the extreme case, it could be asked that only ‘raw materials’–molecules and structures also found outside of the cell, should be included into this hypothetical pipeline. If successful, the system obtained by this method could be claimed to be a ‘completely artificial life form’–even if the design of genomes and other cell elements was guided by templates from living organisms. In [27] the authors underline the fact that alien prokaryotic genomes fail to give ‘instructions’ to a eukaryotic cell, even when the alien genome is faithfully replicated.

So far, genome synthesis and transplantation in prokaryotes as well as design and construction of chromosomes in eukaryotes, have shown that it is possible to substitute native DNA by artificial sequences, designed using as template the original genome. Currently, these wet-lab models allow the segregation of essential versus ‘unnecessary’ or dispensable genome elements, and are leading to a deeper understanding of the function of each element in the cell.

An application of the knowledge about essential cell components is the construction of synthetic cells. This approach explicitly recognizes the obvious fact that not only genomic elements are needed for cell survival, removing in part the bias towards genomic elements. This ‘bottom-up’ approach [28, 29], includes the synthesis of “protocells”, which are compartmentalized assemblies based on different bio-molecules, from oleic acid vesicles containing elements for RNA replication [30], to protein nano-conjugates including stimulus-responsive biomimetic protocells [31]. From the applied perspective, the concept of a “chassis” cell [32] designed for biotechnological uses has also stimulated research regarding the minimal cell [8].

Between synthetic (‘bottom-up’) and analytical (‘top-down’) approaches for the determination of the essential components necessary to produce minimal cells (see Fig 1 in [8]), we have more integrative means to define these elements from both experimental [33] and computational modeling [34] perspectives.

**Fig 1.**
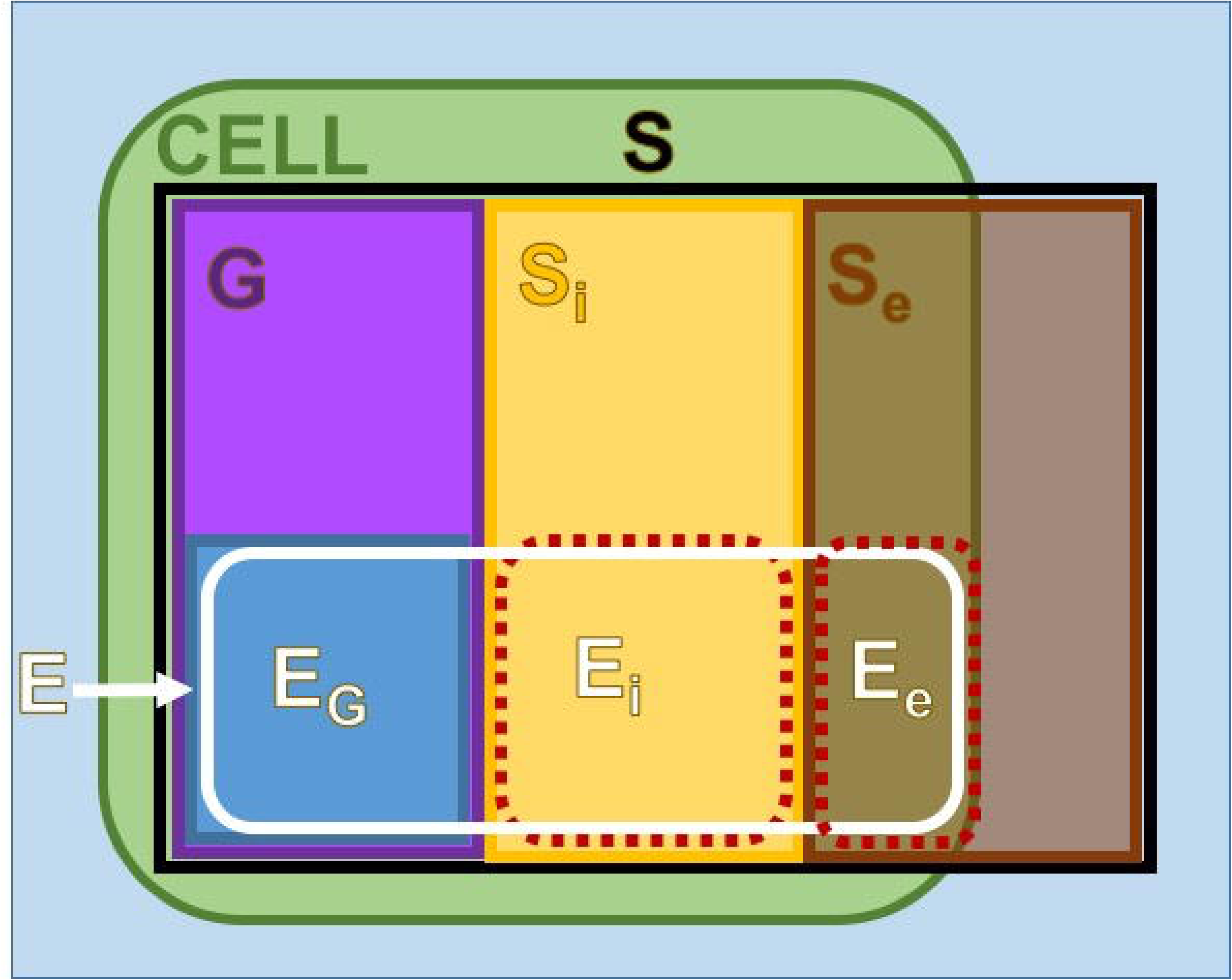
Venn diagram with main subsets of elements in S. **G** - Genome. **S_i_** - Internal elements. **S_e_** - External elements. **E** - Essential elements. **E_G_** - Essential elements within the Genome (the ‘Minimal Genome’). **E_i_** - Essential internal elements. **E_e_** - Essential external elements.

### Theoretical approaches

Whole-cell simulation had been described as a grand challenge of the 21st century [35], asserting that cell behavior cannot be determined or predicted unless a computer model is constructed and computer simulation undertaken. Also, a forum titled “Why build whole-cell models?” [36] underlines the need for data integration for cell modeling and mentions that this integration allows the identification of our knowledge for a given biological system, highlighting poorly understood cellular functions and suggesting areas of research.

E-CELL (http://www.e-cell.org/) was the first software environment for whole-cell simulation initially using *Mycoplasma genitalium* [37]. This platform allows the user to define distinct features of cellular metabolism as a set of reaction rules, and then integrates the differential equations implicitly described in those rules, and shows as a result the dynamic changes in concentrations of cell compounds. The E-CELL environment has resulted in many dynamic simulations of cell processes (see http://www.e-cell.org/publications/).

As mentioned in [34], the main limitations for the construction of whole-cell computational models are the incomplete knowledge of all interactions at the molecular level within the cell, and the fact that no single computational method appears to be sufficient to explain complex phenotypes in terms of molecular components and their interactions. To address those limitations a group lead by Markus W. Covert from Stanford University included all known molecular components and interactions for the life cycle of *Mycoplasma genitalium* into a whole-cell model [34]. They implemented 16 state variables and 28 cell process sub-models, each one analogous to a differential equation, thus the whole-cell model is similar to a system of ordinary differential equations. After initialization of the state variables, the changes of cell state are calculated at temporal steps of one second, allocating and executing cell state variables among sub-models and updating concurrently the values of the states. Each simulation ends when the cell divides or time reaches a maximum value. This approach gave insights into previously unobserved cellular behaviors, such as the rates of protein-DNA association and the inverse relationship between the durations of DNA replication initiation and replication [34].

Other approach to predict essential genes includes the integration of network topology, cellular localization and biological process information [38].

As detailed knowledge about cell components and their interactions increases, that knowledge can be integrated into cell computer models from which emergent –and sometimes surprising, behaviors can arise. *In silico* predictions can then be experimentally tested and more knowledge integrated into the models, leading to a cycle of increased insights into cell function. However, without a solid theoretical foundation for the models, biology will depend only on empirical approximations to understand the phenomenon of life. It is therefore desirable to develop general formalizations of biological principles, which will lead to a more solid philosophical and theoretical framework.

### Binding between molecules: The interactome

Binding between molecules is at the core of biosynthesis. Chemical recognition between proteins, nucleic acids, carbohydrates, lipids and other molecules, drives not only metabolic pathways, but also the assembly of protein [39], ribosomal [40, 41] and transcriptional complexes [42], etc. Molecular recognition and binding has been molded by evolution, resulting in specialization in particular taxa [43].

The ‘interactome’, a term originally coined by Bernard Jacq and co-workers in [44], was defined as ‘*the complete repertoire of interactions potentially encoded by* (*the*) *genome*’. This group underlined the fact that the complexity of an organism is given more by the number of interactions that happen between cell elements than by the number of genes that the organism has. The broad definition of interactome has been delimited to particular types of structures, for example interactions protein–protein [45], RNA-protein [46], RNA-chromatin [47], etc.

Help to study interactomes has come from graph theory [48], which allows a formal treatment of the implicit relations between structures and grants the construction and visualization of biological networks [49, 50] (see also [51] and references thereafter). For example, biological networks based in interactomes have been shown to be useful to identify and study new cellular functions [52], host-microbiota interactions [53], protein communities in addition to disease [45], metabolic [54], motion, adaptive and transport networks [11]. Also, very important theoretical advances in graph theory have been achieved by the study of biological systems [55].

To the best of our knowledge, currently we lack a fully comprehensive interactome, which includes all interactions that could happen between molecules in a given cell. However, this lack of complete knowledge does not preclude fruitful theoretical research to gain knowledge about biological systems, by using current information and making reasonable assumptions. Based on this framework, we assume that there is at least a partial knowledge of the cell interactome, and demonstrate how an algorithmic approach can distinguish essential from non-essential elements.

Given that experimental approaches to determine essential cell components rely in negative results, e.g., a cell in which a gene that codes for an essential structure is disrupted will not survive, we propose that the best method to determine essential cell components is to use properties of the synthesis interactome.

Any mathematical model must disregard some aspects of the phenomenon being modeled, while abstracting the most relevant features and their relations into analytical formulae [56]. Here we present a very general but simplified framework for the different elements and their relationships in an idealized living cell, concentrating into the synthesis of components as function of their binding, and ignoring all complications given by energy transfer, compartmentalization, concentrations and a very long list of etceteras. This scheme, by its simplicity, allows us to show and comprehend how essential cell elements can be distinguished from the nonessential.

## Results and Discussion

### A simple framework for cell elements

Assume that we can make a list of all distinct elements that could exist in a bacteria, within the period immediately after division, that is ‘cell birth’, and just before the initiation of DNA replication–known as the “B period” [57]. With the word ‘element’ in the previous sentence, we refer to components of the cell which form a stable molecular entity, ranging from simple compounds taken from the cell’s environment, metabolites produced inside the cell, to complex molecular arrangements such as membranes, proteins and ribosomes. Fig 1 presents this set as ‘**S**’ and divides it into three disjointed sets, the genome, ‘**G**’, the set of elements synthesized inside the cell, ‘**S_i_**’, and the set of external elements, ‘**S_e_**’. Note that members of this last group can also exist outside the cell, as indicated in the figure. Inside each one of the **G**, **S_i_** and **S_e_** sets of Fig 1 we define subsets ‘**E_G_**’, ‘**E_i_**’ and ‘**E_e_**’, respectively, which represent essential cell elements, i.e., molecular entities without which the cell will be unable to survive and reproduce. As shown in Fig 1 the union of **E_G_**, **E_i_** and **E_e_** is defined as the set of essential elements, **E** = **E_G_** ∪ **E_i_** ∪ **E_e_**. The objective of this notation is to show that, if some assumptions are accepted, it is possible to algorithmically define the set of essential elements, **E**, as function only of the interactions between elements and characteristics of the formulae to synthesize them.

For our aims, the genome can be simply defined as the set of unique DNA sequences that exist within the cell, i.e., if two or more copies of the same DNA molecule exist, then they count only once. It can be rightly argued that the set of chromosomes belongs to the collection of molecules synthesized within the cell, and thus they belong to the set **S_i_**. However, note that we are taking as time window for this model the B period, immediately after division and just before the initiation of DNA replication [57] and during this time period the genome is relatively static within the cell, except possibly for changes in physical conformation, methylation and association with particular structures, but, importantly, we will assume that the sequences of bases of the chromosomes do not change within the B period, and this justifies the separation of the genome, the **G** set, from the set of internal elements, **S_i_**. A more convenient way to define the genome within our framework is as the set of all sub-strings of the bases {*A, T, G, C*} that exist within one or more of the chromosomes of the cell. With such definition **G** is a very large set of DNA strings with sizes 1, 2, …, *r*; where *r* is the size in base pairs of the largest chromosome.

### Binding between pairs of elements and the interactome

#### Binary operators

As mentioned in the introduction, synthesis of cell components depends on binding between molecules. Here we represent the binding between pairs of elements as binary operators of the form

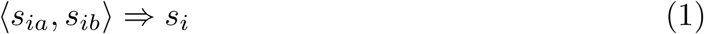

where the operands, *s*_*ia*_ and *s*_*ib*_, are elements that belong to the set **S**, while *s*_*i*_ is a member of the elements synthesized within the cell, **S_i_** (see Fig 1). The operator, “〈, 〉” implies that when the elements represented by the operands are in close proximity under particular conditions, this results into the instantaneous synthesis of the element *s*_*i*_, and we will agree in that the binding operation is commutative, i.e., the order in which the operands are presented is not important and thus 〈*a, b〉* = 〈*b, a* 〉.

The expression (1) for a binary operator is able to represent in an unified way any binding between cell elements, as for example protein–protein interactions [58], protein-DNA bindings [59], and even the status of light sensing proteins [60], before and after receiving the stimulus, etc.

The selection of binary operators to present a general framework for the synthesis of cell structures obeys the fact that interactions of higher order, as for example, bindings between three, four, or more elements, can always be represented as strings of binary operators. Therefore if we assume that three cell elements, say *a, b* and *c*, have binding affinities such that they will form the new structure, *d* = [*abc*], then the synthesis of element *d* can be represented by a string of two nested binary operators, for example as

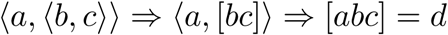

or by other binding order (for example 〈*c*, 〈a, b〉 〉), if order is important. This form of representation by binary operators is then completely general for elements that are synthesized from an arbitrary number of original units.

As an example of the algebraic approach to represent the synthesis of a cellular element let’s take the enzyme RNA polymerase, which will be abbreviated here as ‘*pol*’. For this simplified illustration we will consider that *pol* is constituted only by *α, β* and *β*′ subunits, ignoring the important *σ* factor [61], but considering the *ω* subunit [62], thus we consider *pol* = 2*αββ*′*ω*, because there are two *α* subunits in this enzyme. Now we can substitute in the expression for *pol* the 2*α* part by the corresponding binary operator 〈*α, α* 〉, because we know that 〈 *α, α* 〉 ⇒ 2*α*, and so on, until we decompose *pol* into their subunits, say

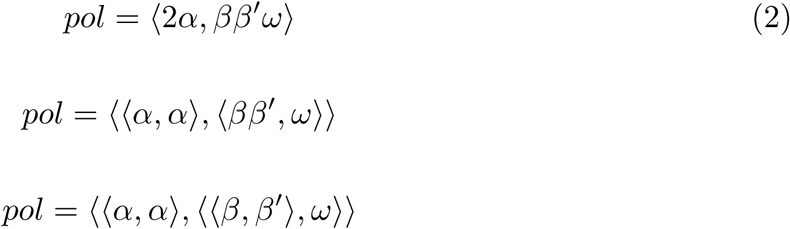

Despite the fact that the synthesis of each *pol* subunit (*α, β, β*′ and *ω*) is complex [63], we can give a more expanded formula for the synthesis of *pol*, expressing the synthesis of each one of its subunits as function of the interaction between the ribosome, ‘*rib*’, and each one of the corresponding transcripts; for example, to synthesize the *α* subunit we need its transcript, say *t.α*, and the ribosome, *rib*; this synthesis is expressed by the binary operator 〈*t.α, rib*〉 ⇒ *α*, and so on for the remaining subunits. By making the corresponding substitutions we find

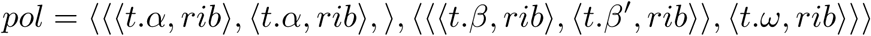

Each one of the transcripts (*t.*) can be expressed by a binary operator involving its gene (*g.*) and, interestingly, the RNA polymerase; e.g., the binary operator 〈*g.α, pol*〉 ⇒ *t.α* showing that to obtain the transcript for the *α* subunit we must have the corresponding gene, *g.α* and *pol*. Finally we obtain an ‘expanded’ formula for *pol* given by

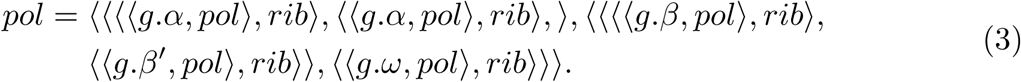

The intriguing fact about Eq (3) giving the ‘expanded’ formula for *pol*, is that it explicitly shows that ‘*to synthesize pol you must have pol*’; i.e., this formula is <u>recursive</u> (or ‘circular’), because it contains between its operands, at the right hand side of the equation, the same term that is being defined, *pol*, at the left hand side of the equation. Even when the fact that to obtain *pol* the cell must have preexistent *pol* molecules is trivially known, the interesting part is that we obtained the expanded formula in (3) from the ‘compact’ form in Eq (2) by a simple ‘recipe’ or ‘algorithm’. Note that if we continue the substitution process in Eq (3) we fall into a never ending loop; on a second round of substitutions to decompose *pol* into their subunits we will have ‘new’ *pol*’s in the formula, and so on. An example of a recursive formula in mathematics is given by the definition of the factorial of a natural number, *n* = 1, 2, 3, as *n*! = *n* ×(*n –* 1)!, together with the agreement that 1! = 1.

Certainly it can be argued that the representation for the synthesis of *pol* in Eq (3) ignores many important facts of the process; for example, for the expression of each gene, the polymerase must recognize a particular motif in the DNA and bind to a particular *σ* factor, say *σ\**, etc. Then, instead of doing the substitution *t.α* = 〈 *g.α, pol*〉 we must expand it to *t.α* = 〈 *g.α,*〈*σ*, pol*, 〉〉 etc. However, to certain extent–which will be discussed later, this ‘lack of detail’ will not affect our conclusions.

#### The Synthesis Interactome (SI) as a list of binary operators

In a first instance we will consider the cell in the period between divisions–the ‘B period’ [57]; later we will examine the phase of DNA replication and mitosis. We also modify the definition given in [44], and consider the Synthesis Interactome (SI) as “*the set of binary operators* (*interactions*) *that result in the synthesis of cellular elements*”. Table 1 presents the scheme for this SI as well as the conditions that must be satisfied by ‘well formed SIs’.

**Table 1.** The Synthesis Interactome (SI).

| Name | Binary operator |
| --- | --- |
| $s_1$ | $\langle s_{1a}, s_{1b} \rangle$ |
| $s_2$ | $\langle s_{2a}, s_{2b} \rangle$ |
| … | … |
| $s_i$ | $\langle s_{ia}, s_{ib} \rangle$ |
| … | … |
| $s_k$ | $\langle s_{ka}, s_{kb} \rangle$ |

Conditions for a well formed SI: i) All represented elements, say

*s*_*i*_, *s*_*ia*_, *s*_*ib*_; *i* = 1, 2, *… k*, must be elements of the set **S** of cell elements (see Fig 1). ii) All names of elements (in column ‘Name’), say *s*_1_, *s*_2_, *…, s*_*k*_, must designate different elements, i.e., *s*_*i*_ ≠ *s*_*j*_ for all pairs *i* ≠ *j*. iii) All *k* binary operators (in column ‘Binary operator’) must be different.

The construction of the interactome for the synthesis of cell elements, or ‘synthesis interactome’ (denoted as ‘SI’; Table 1, as well as in the remaining text), represents the bare minimum to give a logical framework for component synthesis. For example, it does not include any ‘instructions’ for degradation or catabolism, and thus represents only one aspect of cell functions. On the other hand, SI as presented in Table 1 grants the possibility of coding the synthesis of any cell component, breaking it down to the simplest representation: binary operators, or ‘condensed’ formula for synthesis.

Let’s now examine the rules to obtain a ‘well formed SI’, given in the foot notes of Table 1. First, in (i) we ask that all elements named in the SI must be ‘cell elements’, i.e., they must exist in the set **S** of Fig 1. Note that this do not implies that such elements must be present constitutively in all cells; the set **S** denote only elements that can potentially exist in the cell. **S** represent our universe of discurse or ‘universal set’. Now in (ii) it is asked that all elements *s*_1_, *s*_2_, …, *s*_*k*_ in column ‘Name’ must be different. This implies that our SI is non-redundant; for any element synthesized exists one and only one row in Table 1. Condition (ii) also defines the set of elements synthesized within the cell, **S_i_**, because for each *s*_*i*_ we have a binary operator that determines its synthesis; thus **S_i_** = {*s*_1_, *s*_2_, *s*_*k*_}. Note that any particular SI does not need to be ‘complete’ in the sense of listing all posible cell elements; in fact, the task of obtaining a complete SI for any particular specie seems formidable, even with the current large quantity of omics data. Cell elements not found in **S_i_** (column ‘Name’ Table 1) must belong to the complement of this set, say 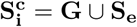 (see Fig 1); i.e, they must be, either, genomic components in **G** or ‘external’ elements in **S_e_**. In a truly complete SI, all elements of **S_e_** must be really ‘external’ to the cell, in the sense of being obtained from the extra-cellular environment; however, in any incomplete SI the set **S_e_** could contain elements which are in fact synthesized within the cell, but for which there is not yet synthesis information in the SI. Finally, condition (iii) implies that there is not any redundancy in SI. In order to observe this assume that there are two rows, say row *i* containing ‘*a*’ in ‘Name’ and ‘〈*b, c* 〉’ in ‘Binary operator’ and a row *j* with ‘*d*’ in ‘Name’ and ‘〈*c, b* 〉’ in ‘Binary operator’. Rows *i* and *j* do not break rule (ii) (because *a* ≠ *d*), however they break rule (iii), because ‘〈*b, c*〉 = 〈*c, b*〉’ (given that binary operators are commutative). The example shows a case where two operands will give different products of synthesis, and this will break the logic scheme of the SI.

Note that all elements listed in the ‘Name’ column of Table 1 belong to the set of internal elements, **S_i_**, while the operands of the binary operators (elements *s*_*ia*_, *s*_*ib*_ in column ‘Binary operators’ of Table 1) are only restricted to be members of **S**.

Further attributes can be added to Table 1 to define, for example, to which particular subset of **S** the operators *s*_*ia*_ and *s*_*ib*_ belong. In the supporting file ‘S1 Text’ we present various examples of SIs, including one which contains information for the synthesis of RNA polymerase, the ribosome and the metabolite streptomycin, while Table 2 presents subset of this SI which include synthesis information only for the RNA polymerase.

**Table 2.** Synthesis Interactome (SI) for RNA polymerase (*pol*).

| Row | Name | Binary Operator | 1st Set | 2nd Set | Type Name | 1st Type | 2nd Type |
| --- | --- | --- | --- | --- | --- | --- | --- |
| 1 | bpb | $\langle bp, b \rangle$ | $S_i$ | $S_i$ | protein complex | peptide | peptide |
| 2 | 2a | $\langle a, a \rangle$ | $S_i$ | $S_i$ | protein complex | peptide | peptide |
| 3 | ba | $\langle bpb, 2a \rangle$ | $S_i$ | $S_i$ | protein complex | protein complex | protein complex |
| 4 | pol | $\langle o, ba \rangle$ | $S_i$ | $S_i$ | enzyme | peptide | protein complex |
| 5 | t.bp | $\langle g.bp, pol \rangle$ | $G$ | $S_i$ | transcript | gene | enzyme |
| 6 | t.b | $\langle g.b, pol \rangle$ | $G$ | $S_i$ | transcript | gene | enzyme |
| 7 | t.a | $\langle g.a, pol \rangle$ | $G$ | $S_i$ | transcript | gene | enzyme |
| 8 | t.o | $\langle g.o, pol \rangle$ | $G$ | $S_i$ | transcript | gene | enzyme |
| 9 | bp | $\langle t.bp, rib \rangle$ | $S_i$ | $S_e$ | peptide | transcript | ribosome |
| 10 | b | $\langle t.b, rib \rangle$ | $S_i$ | $S_e$ | peptide | transcript | ribosome |
| 11 | a | $\langle t.a, rib \rangle$ | $S_i$ | $S_e$ | peptide | transcript | ribosome |
| 12 | o | $\langle t.o, rib \rangle$ | $S_i$ | $S_e$ | peptide | transcript | ribosome |

Keys for element names: ‘pol’ = RNA polymerase, ‘rib’ = Ribosome, ‘a’ = $\alpha$ , ‘b’ = $\beta$ , ‘bp’ = $\beta'$ , ‘o’ = $\omega$ , ‘2a’ = $2\alpha$ , ‘bpb’ = $\beta\beta'$ , ‘ba’ = $2\alpha\beta\beta'$ . Gene names begin with ‘g.’ while transcript names begin with ‘t.’. Columns ‘1st Set’ and ‘2nd Set’ give the sets in which the first and second operands of ‘Binary operator’ exist. Columns ‘Type Name’, ‘1st Type’ and ‘2nd Type’ give the types of elements for column ‘Name’, and the first and second operands of ‘Binary operator’, respectively.

In Table 2, apart from the core columns that determine the SI, say ‘Name’ and ‘Binary operator’ (see Table 1), we included auxiliary columns to indicate to which sets the first and second operands of the binary operators belong, as well as columns giving the type of element of each one of the operands. For technical reasons the Greek letters denoting the subunits of the RNA polymerase were substituted by latin characters.

From Table 2 we can extract the information about members of each subset of **S**, say, the elements which synthesis is defined in the SI:

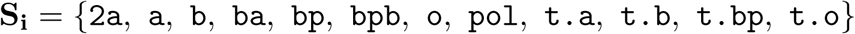

the ones belonging to the genome:

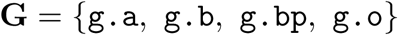

and note that the only element which is not defined within this SI, and thus is cataloged as ‘external’, is the ribosome:

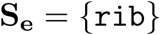

We can say that the SI presented in Table 2 for the RNA polymerase is ‘rooted’ at the ribosome, meaning that this element is not defined within this SI. However, more rows can be added to Table 2 in order define the synthesis of the ribosome; in fact in ‘S1 Text’ we present a more complete SI that includes such information.

### From ‘Condensed’ to ‘Expanded’ expressions: The ‘C2E’ algorithm

As shown in the previous section, the construction of an SI from the core of binding affinities between components, which result in the synthesis of more complex elements, can be achieved by adding knowledge about the behavior of cell components, and in principle this can be automatically accomplished by querying existent databases. For example, the ENCODE (Encyclopedia of DNA Elements) project [64], is building a comprehensive list of DNA motifs which are bound by transcription factors, while the ‘Interactome Projects at CCSB’ [65] are obtaining extensive protein–protein interactome data, etc. However, information in an SI as defined in Table 1 and exemplified in the previous section (Table 2), do not explicitly allow decisions to be made about the essentiality of a cell structure. To do this it is necessary to algebraically ‘expand’ the ‘condensed’ synthesis formula given as a binary operator in the SI. The algorithm to obtain an expanded from a condensed formula (named ‘C2E’) is commented in the ‘**Methods**’ section, and its definition, implementation and practical use are given in ‘S1 Text’, together with the results of applying C2E to the RNA polymerase ‘`pol`’.

By inspecting all the formulae resulting from applying C2E to `pol` in ‘S1 Text’, we confirm that for all `pol`’s components, the corresponding expanded formulae are recursive, i.e., in all cases the formula for the element being synthesized contains within its operands the element being defined.

To give examples of formulae that are not recursive, we present the synthesis of streptomycin, a secondary metabolite exhibiting antibiotic activities, and which is produced by bacteria in the in the genus *Streptomyces* [66]. The SI for streptomycin synthesis was summarized from [67], and the results of applying the C2E to this SI are presented in ‘S1 Text’. These results show that all expanded formulae for each one of the components of this antibiotic, as well as for streptomycin itself are none-recursive, i.e., ‘*to synthesize streptomycin the cell do not need preexistent molecules of streptomycin*’. This is in contrast with the case of the RNA polymerase, where all expanded formulae for each one of the components as well as for the full enzyme were recursive.

### Recursion and essentiality

Assume that we detect an internally synthesized cell element, say *x*, and also independently conclude that to synthesize *x* the cell must have preexistent *x*. This means that the mentioned element, *x*, has a recursive formula and this fact is the way in which we axiomatically define the essentiality of a cell component.

It is practically impossible to experimentally confirm, in every possible case, the fact that recursive elements are indeed essential for the cell. That will entail to be able to eliminate from the cell every representative of the element in question and observe that this causes cell death. However, the logical foundation for this definition of essentiality of a cell element is: 1) We observe a cell element *x* which we know is internally synthesized; 2) We confirm that to synthesize *x* the cell must have pre-existence of *x*, i.e., *x* has a recursive formula. Then we conclude that *x* must be always be present at the cell, at all states of development and at all times. Otherwise, the presence of *x* in the cell is inexplicable, given that *x* is internally synthesized

We agree in that the causal link between our definition of essentiality of cell elements and experimentally testable cell essentiality is subtle; however, as in Physics, we can perform ‘mental experiments’. All biologists will admit that if every molecule of RNA polymerase is eliminated from a cell–without affecting any other cell component, that cell will inevitably die. And the same will happen if the elements eliminated are, for example, ribosomes, or in fact any other ‘essential’ elements. At each one of these putative cases particular arguments can be wield; for RNA polymerase it can be said, ‘*the impossibility to perform transcription will cause a total cell arrest and eventually death*’, and similar statements for other cases. Examples of essential internally synthesized elements are given by the components of the translation machinery [68] for all cell types, actin for eukaryotic cells [69], etc.

On the other hand, let’s examine the negation of our essentiality definition by saying ‘*an internally synthesized element x is essential for the cell, however the formula for the synthesis of x is non recursive*’. We can immediately see that this statement is contradictory, because if the formula for *x* is non recursive, that means that *x* can be synthesized from other cell components, all of them different to *x* and thus *x* could not be ‘essential’–it could be synthesized from a set of elements which essentiality is not known *a priori*.

From a logical point of view we have seen that the fact that an internally synthesized structure *x* has a recursive formula is a necessary condition for *x* to be essential.

In the previous section we have seen that using the information of an SI we can obtain expanded formulae for the elements which synthesis is described in the SI (the elements in **S_i_**), and how in some case these expanded formulae are recursive while in others they are not.

We have exemplified the expansion of formulae for cell elements, but there are cases where such formulae are not ‘closed’, and the substitution process can go on endlessly, increasing the number of operands at each step. Nonetheless, the number of distinct operands that enter into a formula is always finite and can be computed (for details please see the ‘**Methods**’). Let’s denote the complete set of operands that exist in a formula for a structure ‘*x*’ as ‘𝒪*\**(*x*)’.

With this notation we can define our first essentiality rule, say

#### Essentiality rule 1 (ER1)

Let *x* be an internal cell element (*x* ∈ **S_i_**) and 𝒪*\**(*x*) be the complete set of operands (elements) that exist into its expanded formula. Then *x* will be essential for cell surviving if

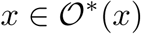

i.e., if *x* is a recursive structure.

The rationale for statement **ER1** resides in the fact that if *x* is recursive, then such element cannot be synthesized ‘*de novo*’ in its absence, e.g., ‘*to synthesize RNA polymerase the cell must have RNA polymerase*’, etc.

One can question if the degree of ‘detail’ embedded into the SI for the synthesis of *x* will affects the validity of **ER1**. In fact, if there is not ‘enough’ information for the synthesis of an structure *x* into an SI, the recursiveness of its formula could not be discovered. For example, we found that the formula for the synthesis of the RNA polymerase, ‘`pol`’, was recursive only when we took into account the transcripts that are needed for the synthesis of its subunits: `1 t.bp`, `t.b`, `t.a` and `t.o` (see Eq 3); if we eliminate from the SI the rows in which those transcripts are defined, we still have a valid SI, which still contains partial information for ‘`pol`’ synthesis, however by analyzing such reduced SI we will not be able to declare ‘`pol`’ as recursive and thus as essential by using **ER1**.

The previous example shows that evidence of essentiality can only be obtained if ‘enough’ information about the synthesis of an element is present in the SI analyzed. *A priory*–without performing calculations, it is difficult to say by observing an SI, if it contains enough information to determine which structures are essential by rule **ER1**. However the algorithm presented in the **Methods** section determines the complete sets 𝒪 *(*s*_*i*_) for all *s*_*i*_ ∈ **S_i_**, allowing the application of **ER1**.

Because at the deepest level the synthesis of any internal cell element depends, directly or indirectly, on the information given by the genome, one can hypothesize that SIs integrating all necessary elements of **G** among its operands (elements *s*_*ia*_, *s*_*ib*_ in column ‘Binary operators’; see Table 1), will give enough information to determine essentiality of the corresponding internal elements. Nevertheless that is not always the case (see ‘S1 Text’ for a counterexample)

On the other hand, an ‘excess’ of detail or information about the synthesis of a given structure could not revert essentiality classification when it has been stablished using **ER1**. For example adding rows to the `pol` SI (S1 Text) to include other genomic and regulatory elements for the expression of ‘`pol`’ will not alter the fact that it will be classified as essential, even if the expanded formula changes, increasing in complexity and an increase is also observed in the number of operands needed for its synthesis.

To complete the set of essential cell elements we present a second rule of essentiality, say

#### Essentiality rule 2 (ER2)

Let *x* be an essential structure which complete set of operands is 𝒪*(*x*). Then all elements of 𝒪*(*x*) are essential.

This rule affirms that all elements that enter into the synthesis of an essential element are also essential (note that *x* ∈𝒪*(*x*), given that *x* fulfills **ER1**). To see the logic of **ER2** note that, given that 𝒪*(*x*) is the complete set of operands to synthesize *x*, each and every one of the elements of 𝒪*(*x*) must be present in the cell for *x* to exist in the cell. Now assume that *x*\* is an element of 𝒪*(*x*), i.e., *x*\*∈𝒪 *(*x*). Then *x*\* is essential, because without it the synthesis of *x* cannot be completed. Assuming that *x*\* is not essential leads to a contradiction, because that will imply that *x* is also not essential, a fact that is not under discussion.

**ER1** defines essentiality for elements synthesized within the cell (in **S_i_**) while **ER2** extends this property to any member of all elements of the cell (**S**), which satisfy the condition to be members of one or more of the sets of complete operands for essential elements. Thus elements that fulfill **ER2** can be members not only of **S_i_**, but also of **G** or **S_e_**, i.e., genomic or external elements. Together **ER1** and **ER2** give necessary (**ER1**) and sufficient (**ER2**) conditions for essentiality of cell elements, allowing to define the set of essential elements shown in Fig 1 as

#### Set of essential elements E

Let **E_i_** = {*e*_1_, *e*_2_, *…, e*_*k*_} be the set of all elements that fulfill **ER1**, i.e., the set of essential structures such that **E_i_** = {*e*_*i*_*|e*_*i*_ ∈𝒪*(*e*_*i*_)} where 𝒪*(*e*_*i*_) is the set of complete operands for *e*_*i*_; *i* = 1, 2, …, *k*. Then the complete set of essential structures, **E**, is given by

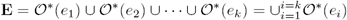

i.e., the set of essential structures is formed by all elements that follow rules **ER1** or **ER2**.

As mentioned in [34], one of the main limitation for the construction of whole cell computational models is the incomplete knowledge of all molecular interactions within the cell, and, as the authors say in [16], ‘*No single cellular system has all of its genes understood in terms of their biological roles.*’–and the same is true for all interactions between molecules in a cell of a particular species. Complete knowledge of all possible interactions between pairs of molecules in the cell of a given specie is a very stringent condition to set for any practical model. Currently, we are far from that exhaustive knowledge, even for the most simple and well-characterized bacterial models. In [70] the authors developed a method to estimate the size of the protein interactome from incomplete data and estimate for example that there are approximately 650,000 protein pair interactions in humans, however only a relatively small set of these interactions have been experimentally corroborated. Thus we must take into account the fact that almost any SI determined will be to some extent incomplete, and thus ponder the consequences of this fact for the classification of the essentiality of cell elements.

The conditions for a ‘well formed’ SI, given in the foot notes of Table 1, imply that if the table SI* represents a well formed SI with *k* rows, then, any subset of *t* rows of SI* (*t < k*; *t ≥* 1) will also fulfill the conditions to be a well formed SI. At the limit, an SI with *t* = 1 row is a (trivial) well formed SI, and it will inevitably give a non recursive formula for the element defined. Take as example the row 2 of Table 2, which define the synthesis of the 2*α* subunit of the RNA polymerase by the binary operator ‘〈*a,* 〉 *⇒ a* 2*a*’ (in columns ‘Binary operator’ and ‘Name’ respectively). In isolation this formula will give the wrong answer to the question of the essentiality of the 2*α* subunit, classifying it as ‘not essential’. Further discussion of this fact is given in ‘S1 Text’.

### The interactome as a biological network

Even when the algebraic criteria **ER1** and **ER2** are together necessary and sufficient to determine essentiality of a cell component during the B period, this approach is not intuitively appealing, mainly because it lacks a graphical representation from which one could directly corroborate the logic of the results. Fortunately we can use elements of graph theory [48, 71] to visualize the relations in the interactome (see ‘S1 Text’ for the formal definition of a graph).

In fact, the interactome defines two graphs, the ‘binding’ relation, implicit in the binary operators ‘〈*s*_*ia*_, *s*_*ia*_〉’ (see Table 1), and the more complex ‘synthesis’ relation, implicating three actors and represented by the complete binary operators ‘〈 *s*_*ia*_, *s*_*ia*_〉 ⇒ *s*_*i*_’ in the interactome. The former defines an undirected graph, while the later defines a directed graph or ‘digraph’ [71]. The binding relation will show a plot in which pairs of binding elements will be united by an undirected edge (see ‘S1 Text’), while the synthesis relation will generate a graph in which elements will be united to each other by directed edges, or ‘arrows’, as shown in Fig 2.

**Fig 2.**
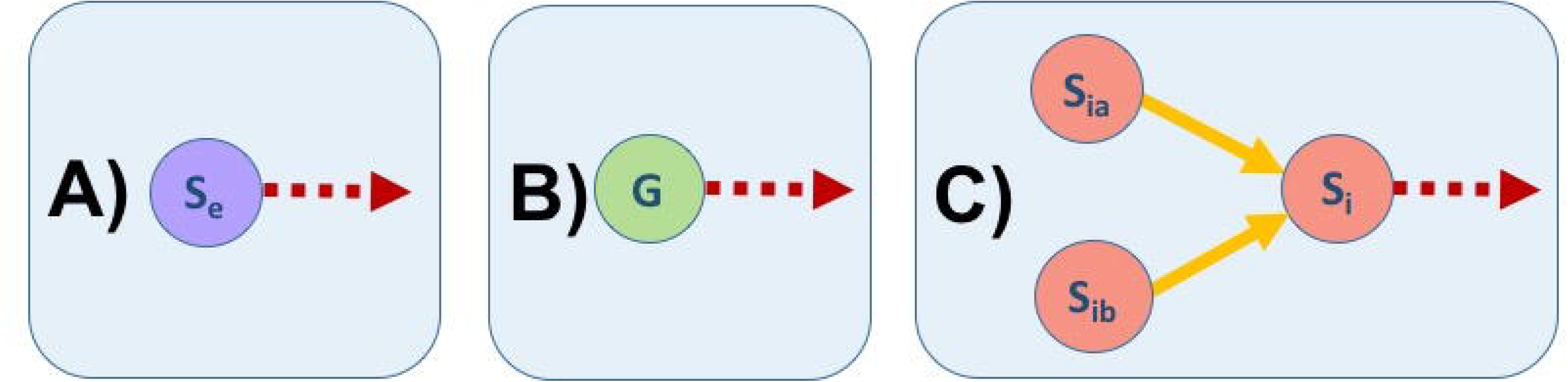
Possible cases for edges in the synthesis interactome (SI) as a network. In (A) and (B), elements of **S_e_** and **G**, respectively, there are only edges directed out of the element (represented by red dashed arrows). In contrast, in (C), for elements of **S_i_** there will be exactly two edges coming from other elements (*S*_*ia*_ and *S*_*ib*_) and there could be any number of edges going out of from *S*_*i*_ to other elements (represented by red dashed arrows).

Fig 2 shows that elements that belong to **S_e_** and **G** (panels **A** and **B** respectively) can only have (one or many) edges that go from the corresponding element to point to other elements. This means that elements in the set of external elements, **S_e_**, or in the genome, **G**, can be used in the synthesis of other elements (the points where the corresponding arrows arrive; not shown), but, there is not information for their syntheses in the SI. On the other hand, internal cell elements in the set **S_i_** must, by the definition of binary operators, be synthesized within the cell by the binding of exactly two elements; that is why there are exactly two arrows arriving to the *S*_*i*_ element (yellow arrows in panel **C** of Fig 2), and there could be one or more arrows departing from *S*_*i*_ (red dashed arrow in panel **C** of Fig 2).

#### Plots of SIs as biological networks

Technical details to study and visualize SIs using the R environment [72] and the ‘igraph’ R package [73] are presented in ‘S1 Text’, while data and functions for interactome study can be downloaded as our R code ‘S1 Binary’. Here we show and comment the results of transforming the SIs presented and discussed above as graphs of biological networks. We will see that the fact that an expanded formula for an element is recursive, implies that such element is part of a ‘closed walk’ [71], i.e., a circle of elements (vertices) and arrows (directed edged) within the graph of the corresponding SI. In other words, synthesis circularity–the need of an element for its own synthesis, is echoed in graph circularity.

Fig 3 shows the biological network resulting from transforming the SI for RNA polymerase (in Table 2) into a directed plot, where vertices (circles) are the elements and directed edges (arrows) give the synthesis relation obtained from the binary operators in column ‘Binary operators’ of Table 2.

**Fig 3.**
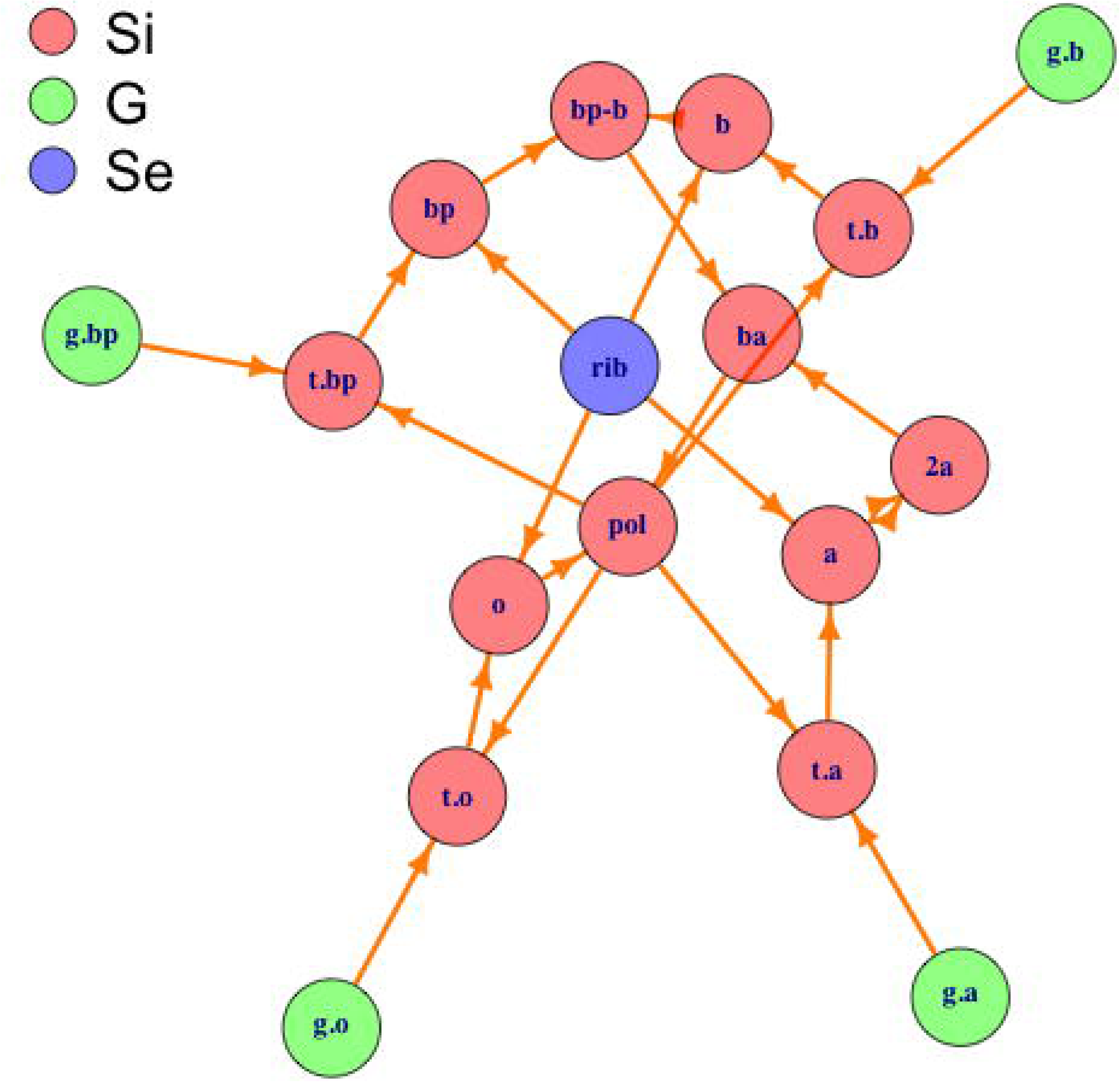
Plot of synthesis SI for RNA polymerase (Table 2) annotated by set of origin. Biological network representation for the synthesis of RNA polymerase colored by set of origin (**S_i_** - Internal elements, **G** - Genes and **S_e_** - External elements). For meaning of the abbreviated element names see Table 2.

In Fig 3 we can see how the synthesis plot of the biological network for RNA polymerase (corresponding to the SI presented in Table 2) shows ‘closed walks’, i.e., cycles that begin and end at each one of the internal elements, **S_i_**, defined by the SI for the RNA polymerase in Table 2. Table 3 explicitly shows each one of these 12 cycles to made it easier to count and follow them in Fig 3.

**Table 3.** Cycles (closed walks) present in the network for RNA polymerase (in Fig 3)

| Name | Cycle |
| --- | --- |
| *2a* | $2a \rightarrow ba \rightarrow pol \rightarrow t.a \rightarrow a \rightarrow 2a$ |
| *a* | $a \rightarrow 2a \rightarrow ba \rightarrow pol \rightarrow t.a \rightarrow a$ |
| *b* | $b \rightarrow bpb \rightarrow ba \rightarrow pol \rightarrow t.b \rightarrow b$ |
| *ba* | $ba \rightarrow pol \rightarrow t.bp \rightarrow bp \rightarrow bpb \rightarrow ba$ |
| *bp* | $bp \rightarrow bpb \rightarrow ba \rightarrow pol \rightarrow t.bp \rightarrow bp$ |
| *bpb* | $bpb \rightarrow ba \rightarrow pol \rightarrow t.bp \rightarrow bp \rightarrow bpb$ |
| *o* | $o \rightarrow pol \rightarrow t.o \rightarrow o$ |
| *pol* | $pol \rightarrow t.bp \rightarrow bp \rightarrow bpb \rightarrow ba \rightarrow pol$ |
| *t.a* | $t.a \rightarrow a \rightarrow 2a \rightarrow ba \rightarrow pol \rightarrow t.a$ |
| *t.b* | $t.b \rightarrow b \rightarrow bpb \rightarrow ba \rightarrow pol \rightarrow t.b$ |
| *t.bp* | $t.bp \rightarrow bp \rightarrow bpb \rightarrow ba \rightarrow pol \rightarrow t.bp$ |
| *t.o* | $t.o \rightarrow o \rightarrow pol \rightarrow t.o$ |

From Fig 3 and Table 3 we can see that there is a correspondence between recursive elements uncovered by the C2E algorithm and closed walks (cycles); in fact, to each internal element that has a recursive formula, corresponds a closed walk in the network; graph theory unveils the essentiality of the elements in a way analogous to the algebraic substitutions performed by C2E. In Fig 3 only external elements in the **S_e_** set, say the genes for the RNA polymerase, *g.a, g.b, g.bp* and *g.o* (shown in the periphery of the network as green circles), and the ribosome, *rib* (at the center; violet circle) are not included into a cycle. As mentioned before, these ‘external’ elements are not defined within the SI, and thus form the ‘root’ of that graph, i.e., the elements from which the synthesis of all the others elements begins. In fact, there are graph theory algorithms to find closed walks for an element within a network [73].

‘Name’ - Name of each one of the elements in the set of internal elements, **S_i_**. ‘Cycle’ - Closed walk beginning and ending at element ‘Name’. Edges (directed arrows) are symbolized as ‘→’.

Fig 4 shows the network resulting from the partial SI for RNA polymerase. This partial SI results from deleting rows 5 to 8 in Table 2; i.e., we deleted all the rows that defined the synthesis of the transcripts (elements which name begins with ‘`t.`’) for each one of the subunits (`a, b, bp, o`) from their corresponding genes (elements which name begins with ‘`g.`’).

**Fig 4.**
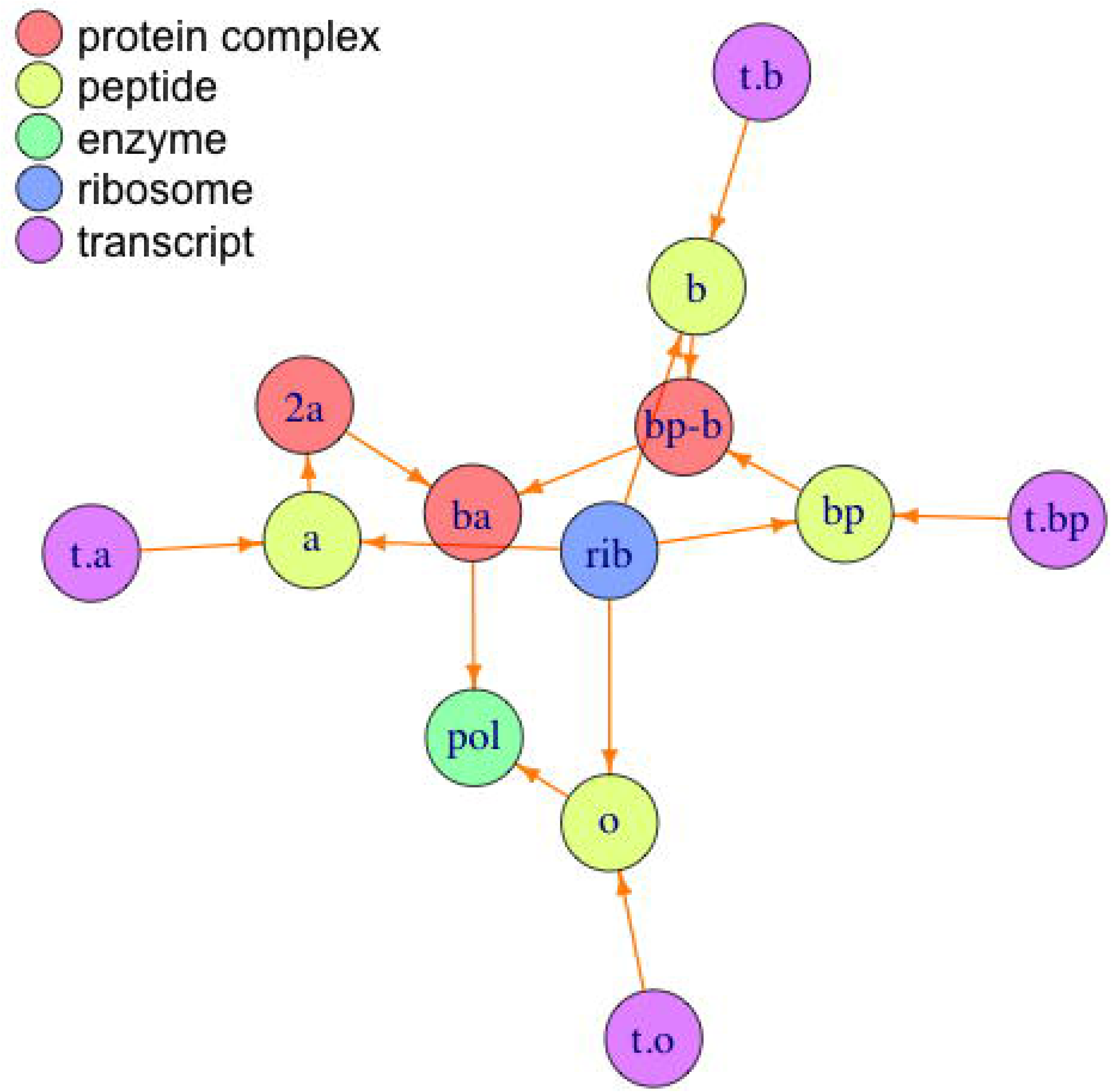
Plot of partial synthesis SI for RNA polymerase annotated by type of element. Biological network representation for the partial synthesis of RNA polymerase colored by type of element. For the meaning of the abbreviated element names see Table 2.

In contrast with Fig 3, Fig 4 do not have any closed walks (cycles) for any of the elements present in the plot. From Fig 4 it can be verified that departing from any one of the elements it is impossible to comeback to the same element, and this is a result of the fact that the synthesis of the components of the RNA polymerase is incompletely described by the corresponding SI. In the partial SI for RNA polymerase the transcripts (elements beginning with ‘*t.*’) will be classified as ‘external elements’, i.e., the information for their synthesis is not included into that partial SI; they have only outgoing, but not incoming arrows (see Fig 2), and thus all cycles for the elements in Fig 4 remain as open paths without forming cycles.

The analysis of the partial SI for RNA polymerase, obtained by erasing rows 5 to 8 in Table 2, give only non essential structures (data not shown), because the recursiveness of all the structures is not present in that partial SI. This is also reflected in Fig 4, where no closed walks are found. Thus, there is a correspondence between the negation of **ER1** and the results obtained with graph theory; when an element is non recursive, there is not a closed walk for that component.

Fig 5 presents the network for the SI for streptomycin (‘`STR`’; see ‘S1 Text’).

**Fig 5.**
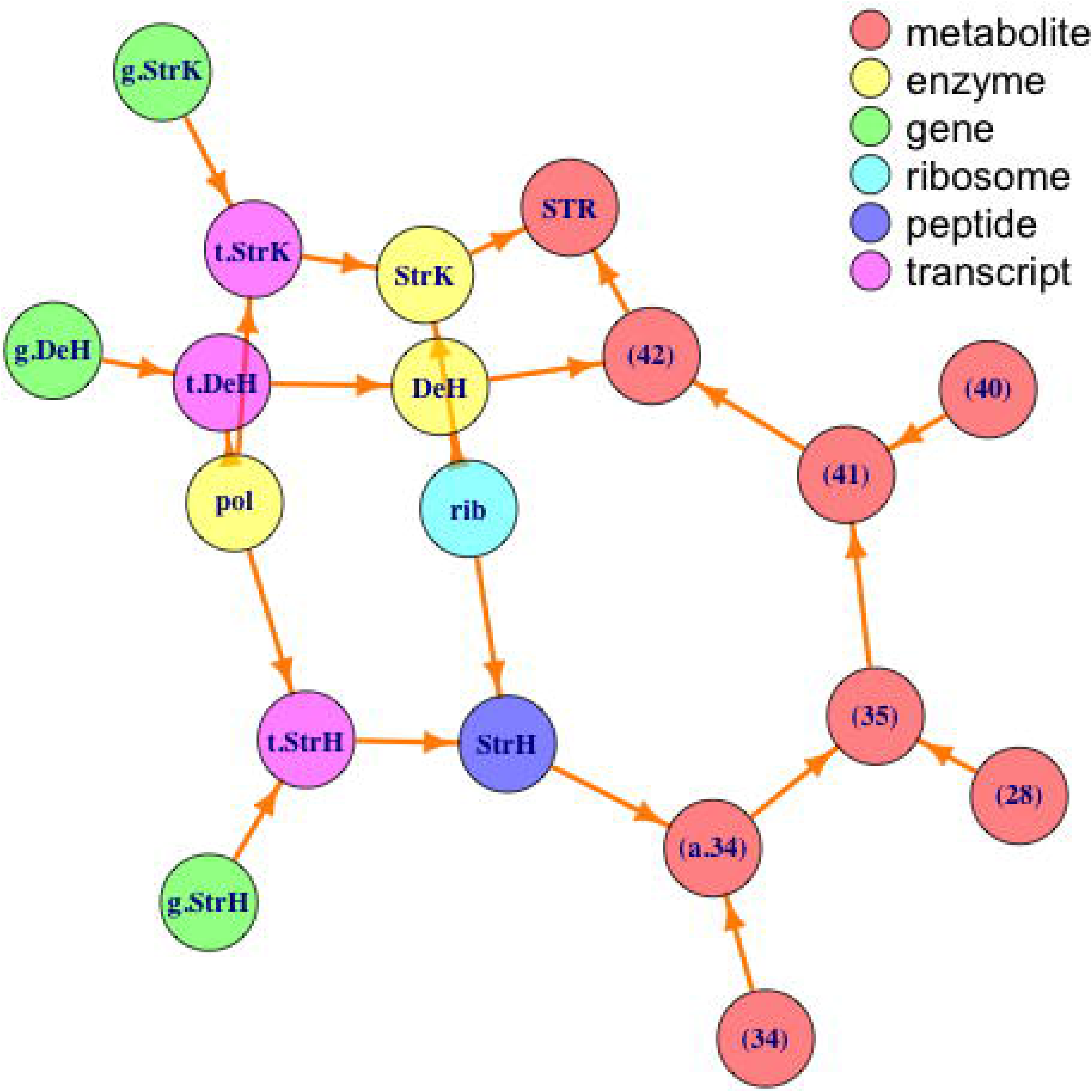
Plot of synthesis SI for streptomycin annotated by type of element. Biological network representation for the synthesis of streptomycin colored by type of element. For the ‘`STR`’ SI and meaning of the abbreviated component names see ‘S1 Text’.

From Fig 5 we can see that there are not closed walks for any of the elements shown, corroborating the result using the C2E algorithm that none of the internal elements whose synthesis is described in the corresponding SI has a recursive formula and, in consequence, non of them is essential for cell survival (see ‘S1 Text’ for details).

Synthesis interactomes (SIs) can be constructed in a progressive manner, by adding rows describing the synthesis of elements which at a previous stage were classified as ‘external’. For example, in the SIs for RNA polymerase (Table 2) and streptomycin (in ‘S1 Text’), the ribosome (*rib*) is considered as an external structure. Nevertheless, by adding rows describing the synthesis of the ribosome from their genes of origin (including the genes for ribosomal RNAs as well as all peptides involved in this structure) we obtain a more ‘integrated’ SI where the synthesis of the ribosome is included. Also, by combining various SIs, without breaking the rules given at the foot notes in Table 1, we can include more elements and ‘details’ about the synthesis of internal elements carried out in the cell. In ‘S1 Text’ and ‘S3 Text’ we present and analyze an integrated SI, which includes the synthesis of RNA polymerase, streptomycin and the ribosome. This procedure can be continued as desired to include more and more elements, until eventually it will include the synthesis of all elements from a given cell species. As an illustration, Fig 6 shows the ‘integrated’ SI including the synthesis of RNA polymerase, streptomycin and the ribosome.

**Fig 6.**
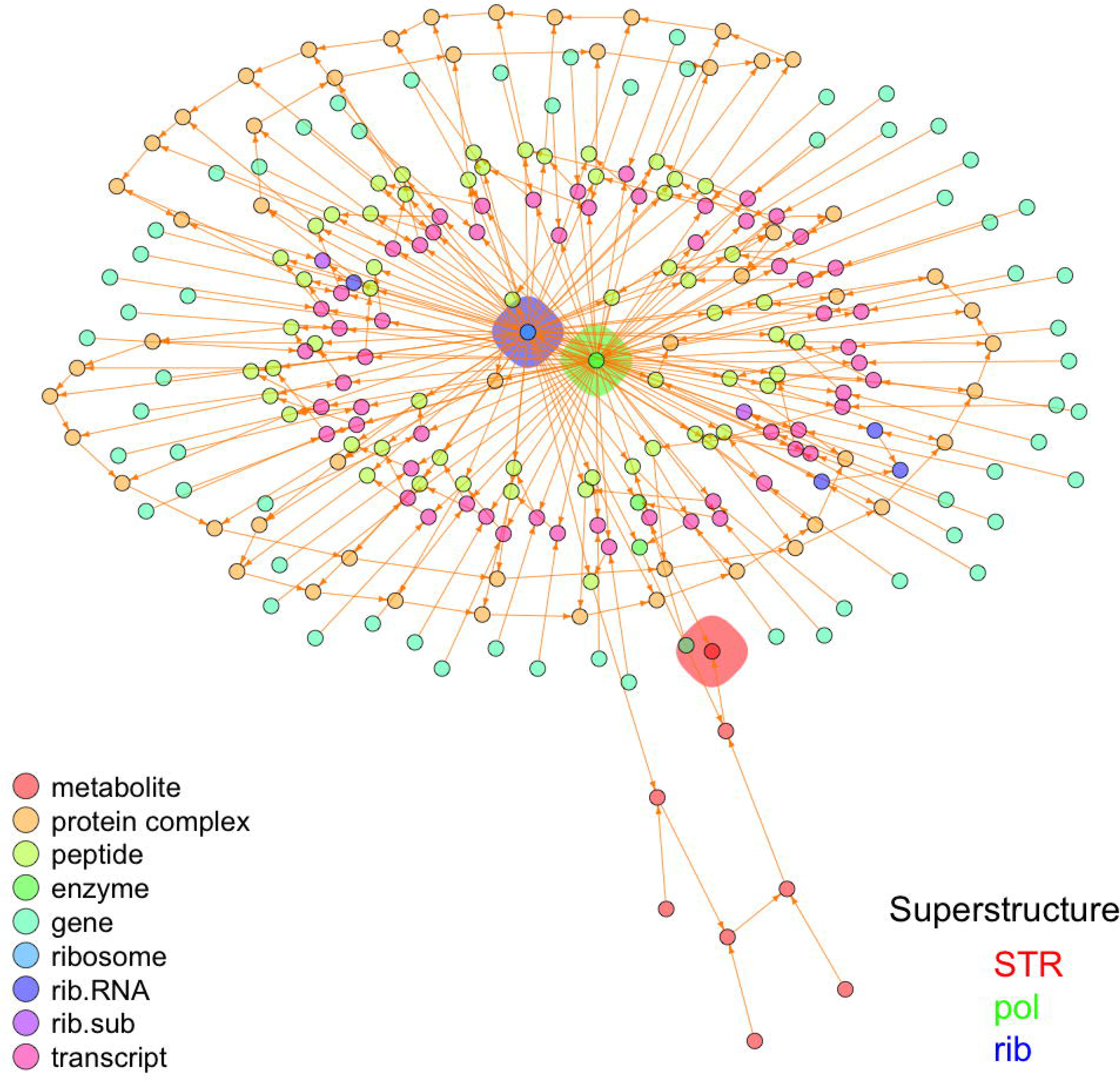
Plot of synthesis SI for the ‘integrated interactome’ annotated by type of element and superstructure. ‘Superstructure’ centers: *STR*-Streptomycin, *pol*-RNA polymerase and *rib*-Ribosome, are annotated by a colored polygon, while elements (circles) are not annotated with labels, but only by type of element in the legend (see ‘S1 Text’).

Fig 6 shows that the ribosome (*rib* in blue polygon) and the RNA polymerase (*pol* in green polygon), are highly connected elements (called ‘hubs’ in the literature–see for example [74]), while streptomycin (*STR*) is not a hub at all, being connected with only two other elements. The fact that both essential elements, *rib* and *pol*, are highly connected hubs, while the secondary metabolite *STR* is not, is in complete agreement with the ‘lethality and centrality hypothesis’ [75] which states that ‘*The most highly connected proteins in the cell are the most important for its survival.*’. In fact, our results allow to expand this hypothesis from ‘protein’ to more general elements (such as the ribosome), and explain in clear terms the essentiality of these hubs by the recursiveness of their expanded formulae, giving an straightforward answer to the question ‘*Why do hubs tend to be essential in protein networks?*’ asked in [76, 77]. Our results also agree with the study of eukaryotic protein-interaction networks [78], where the authors show that proteins with a more central position in the networks are more likely to be essential for survival, regardless of the number of direct interactors. In fact, peptides which form parts of the RNA polymerase and the ribosome form an inner ring in Fig 6.

It is important to underline that in the analysis of the ‘integrated SI’, which defines the synthesis of the secondary metabolite streptomycin (`STR`), the RNA polymerase (`pol`) and the ribosome (`rib`) in a single SI, our algorithmic approach correctly indicates the essentiality of the RNA polymerase and all its components, as well as the essentiality of the ribosome and all its components, but also correctly classifies the secondary metabolite streptomycin and all its components as non-essential cell elements (see ‘S1 Text’ for full results and discussion).

### Essentiality of the genome duplication machinery

Since the year 1858, when R. Virchow expressed his now famous quote, ‘*omnis cellula a cellula*’ [79], it has been completely clear that one of the main attributes of life is cell reproduction, which implies DNA replication. Genomic replication requires a large collection of proteins properly assembled, which are named ‘replisome’ [80]. However, up to this point we have defined cell elements that are essential only during the “B period” [57], i.e., after the end of mitosis and before DNA replication. Without further details, we can close this gap in our definition of the essential cell elements with a third and last rule for essentiality

#### Essentiality rule 3 (ER3): Essentiality of genome replication machinery

Let *g*\* be a genomic element, *g*\* ∈ **G**. Then *g*\* will be essential for genomic replication if by deleting all copies of *g*\* genome replication is impossible.

In contrast with rules **ER1** and **ER2**, **ER3** is not algorithmic, but experimental. The reason for this is that until the DNA replication begins, genes and elements involved with genome duplication can be damaged–for example by mutation, but that damage will be overlooked until the signals for entering into mitosis are sensed [81]; at that point the damage will be evident if genome replication halts. For example, using a gene knockout method in *Halobacterium* the authors in [2] showed that only ten out of nineteen eukaryotic-type DNA replication genes are essential for that bacteria. Those genes code for two of ten Orc/Cdc6 proteins, two out of three DNA polymerases, the MCM helicase, two DNA primase subunits, the DNA polymerase sliding clamp, and the flap endonuclease.

The reason by which **ER3** is not written algorithmically, is that the essentiality of the genome replication machinery is of ‘second order’, in the sense that essentiality is only evident for ‘the next cell generation’. If we include the synthesis of DNA polymerase into an SI (data not shown), the expanded formula for that element do not show recursion, i.e., ‘to synthesize DNA polymerase the cell does not need DNA polymerase’. However, that is true only immediately–in a ‘first order’ sense, because evidently to form DNA polymerase the cell must have come from a (parent) cell that was able to replicate its genome and, obviously, that cell must have had DNA polymerase. To discover the elements determined by **ER3** we need experimental approaches, as for example the ones described in [2, 4, 5, 82, 83].

### The ‘Minimal Set of Preexistent Elements’ (MSPE)

In Fig 1 we show the Venn digram for all cell elements, **S**, which is divided into the disjoint sets of genomic (**G**), internal (**S_i_**) and external elements (**S_e_**),

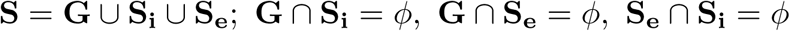

in which *ϕ* denotes the empty set. Also in Fig 1 we show the proper subset of essential elements, **E** ⊂ **S**, which in turn was conceptualized as formed by the essential elements existent in **G**, **S_i_** and **S_e_**, say **E_G_**, **E_i_** and **E_e_**, respectively,

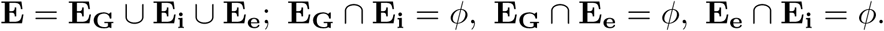

We were able to algorithmically determine all elements of the set of essential internal elements (**E_i_**) by using our **ER1**, which can be restated by saying that all essential elements are ‘preexistent’, because to synthesize any of them they must exist prior to the beginning of the synthesis operation. Later, and using **ER2**, we showed that all genomic or external elements included as operands in the formulae for essential elements were also essential, determining the set **E_G_** ∪**E_e_**. Finally the preexistence (in the previous generation) of the genome replication machinery allowed us to state **ER3**, completing the set **E_G_** with genes that encode for such machinery. *A priori* only **ER3** explicitly demands ‘extra’ experimental work; the other two essentiality rules rely on knowledge about the synthesis of elements in the form of an SI, which for many elements is well characterized and can be obtained from specialized databases and the literature.

Given that, as shown here, ‘preexistence’ of cell elements is the core of essentiality, we propose that the set of essential cell elements could be designated as the ‘Minimal Set of Preexistent Elements’ (MSPE). With the approach presented here, and summarized in **ER1** and **ER2**, it is possible to integrate the information existent about biological synthesis into an increasingly detailed SI for particular species, or in general for full taxa. From such SIs, and by employing **ER1** and **ER2** and the associated algorithms (see ‘**Methods**’ and supporting information), it is then possible to distinguish the majority of the members of the MSPE. In principle, the only elements of the MSPE that will be missed by this approach will be the ones needed for genome replication, which are relatively well known for many organisms (see for example [2]).

Current knowledge about DNA motifs and their interaction with other elements [64], as well as particular interactomes, for example between proteins [84], RNA and chromatin [47], and biochemical networks [47, 85, 86], among others, can be included into SIs to extract the members of the MSPE.

Here we centered in the essentiality of cell elements; however, survival and reproduction of whole multicellular organisms was not discussed. It appear obvious that the set of essential elements at the organism level must be larger than the MSPE that we have presented, as it is evident from the proportion of essential genes at different taxonomical levels [13], discussed in the introduction. In fact, many lethal or detrimental mutations in humans are only evident in infants [87] or even adults [88]. It appears unlikely that the straightforward criteria employed here to define the MSPE could be escalated to fully determine the MSPE for multicellular organisms, given the complex associations implicit in the in the synthesis of multicellular structures such as tissues, organs, etc. However, it is possible that the criterion of circular dependence or recursiveness could be employed with that aim.

### Modifying the SI definition

Conditions for a well formed SI, presented in Table 1 and discussed below in the **Methods** section, were set to show the rationale of **ER1** and **ER2** and facilitate the descriptions of the algorithm to find essential structures. However it is clear that real SIs will not always comply with such conditions. Here we briefly discuss how the relaxation of such assumptions could affect the results presented and which additions could be done to our SI definition to make it more realistic.

Biological networks could be redundant [89] and are in general robust [90]. In contrast, our SI model as defined in Table 1 is non redundant (by condition ‘ii’), and as we have seen non robust, in the sense that the elimination of rows implies differences in the discovery of essential structures. In fact, lack of robustness is in part due to the non redundancy imposed by condition ‘ii’.

Relaxing condition ‘ii’ in Table 1, allowing different binary operators to result in the synthesis of the same external element will produce alternative synthesis pathways for the same element, something that is common in metabolic pathways [91]. Relaxation of ‘ii’ to allow multiple synthesis pathways for the same structure complicates the finding of essential structures–because multiple options need to be taken into account, but does not contravene **ER1** or **ER2**. By modifying ‘ii’ we will have more realistic and robust SIs, complicating computations but without violating essentiality rules.

A more intriguing situation arises if we want to modify ‘iii’ which states that ‘*All k binary operators must be different* ‘. If we allow duplicity (or multiplicity) of binary operations, for example, say that we want to model a case where ‘〈*a, b*〉 ⇒ *c*’ OR ‘〈 *a, b*〉 ⇒ *d*’, i.e., the case where two operands give different products, the only possible solution that we could see is to use stochastic assignation of the result. For example, to choose ‘〈*a, b*〉 ⇒ *c*’ with probability *p* and ‘〈*a, b*〉 ⇒ *d*’ with probability 1 – *p*, etc. At this point it is not clear if such possibility is biologically relevant.

Other aspect in which our SI definition could be developed is the inclusion of time in the model. Definition of our binary operators assume an atemporal model, in the sense that we assume that synthesis interactions are performed ‘instantly’. If we want to include time in the model, we could select discrete intervals and, in the simplest case uniform discrete times for all binary operators. Such modification will give dynamical models, which could be very important for some applications but which will not modify the rules of essentiality.

Multiple possibilities exist to modify the definition of an SI to allow more realistic cases, which will give more precise results than the simple model presented here. In all cases the importance of these models (the one presented here as well as putative modifications) is that in all cases different sources of data must be integrated to model *synthesis* of elements, i.e., it is not sufficient to have isolated interactomes, as protein–protein, DNA-protein, etc.; the synthesis of elements must be completely described in a single and connected SI, because as we have seen only when relatively complete information about the synthesis of a given element is present in the interactome it is possible to decide about it’s essentiality.

### Obtaining the elements of a minimal cell

We have presented an algorithmic definition that allows the separation of essential from dispensable cell elements. To obtain the elements of a minimal cell from the complete SI for that cell specie, it is sufficient to selectively delete the rows of that SI which are exclusively involved with the synthesis of non essential elements–after its determination has been performed using the rules proposed here. Then the practical problem is to obtain such complete SI.

For example, even when *E. coli* is one of the best understood and most analyzed organisms [92], having the best electronically-encoded regulatory network of any free-living organism [93], to the best of our knowledge we currently lack the integration of all this knowledge into a platform focus in the <u>synthesis</u> of the *E. coli* cell elements, fulfilling the model presented here or an improved version of it.

Already the reduction of *E. coli* genome by making precise deletions of non essential genes and sequences has led to unanticipated cell properties [92]; thus we expect that the integration of complete SIs in which our method could identify essential cell elements will advance the understanding of core cell elements and functions.

## Conclusion

Essential cell elements are determined by the fact that their synthesis needs their preexistence. This criterion allows to distinguish essential from non-essential elements in an algorithmic way when enough information is available.

A first question that arises here is which quantity of information is enough to determine essentially of a cell element within an SI using our algorithmic approach. As seen in the example presented for the RNA polymerase, essentiality of the ribosome cannot be judged within the RNA polymerase SI, because there the ribosome is given as an ‘external element’ in **S_e_**; i.e., there is no information for the synthesis of the ribosome in that SI. In contrast, in the integrated interactome (see Fig 6) essentiality of the ribosome can be determined because in that SI ribosome synthesis is defined by binary operators. This can be generalized to say that essentiality of a cell element can be algorithmically decided only when its synthesis is defined, as a set of binary operators, within the corresponding SI. In a complete SI for a given specie, the synthesis of all cell elements must be defined by a set of binary operators, and external elements, **S_e_**, must contain only genomic elements and truly external elements that the cell could obtain from its immediate environment. In contrast with our approach, experimental approaches to determine essential elements rely on negative results (cell inviability) when mutating the genes that determine such elements. Examples are found in [82] for *Bacillus subtilis* and in [94] for *E. coli*. In this last publication the authors were unable to disrupt 303 genes, including 37 of unknown function, which they label as candidates for essential genes.

A second question concerns the complexity and size of a complete SI. As defined here, SIs include as subsets other particular interactomes, as protein–protein, protein-DNA, etc. A relevant question is how large a complete SI of a particular specie will be, and thus how complex is the algorithmic solution that we propose to determine essentiality. We presented an SI (`int.SI`, see ‘S1 Text’ and Fig 6) with 184 binary operators, which includes the synthesis of the ribosome, the RNA polymerase and the antibiotic streptomycin. In this SI the ratio of the number of binary operators to genes included in the SI is 184*/*62 = 2.9677 *≈* 3. Making a linear extrapolation, we could estimate the minimum number of binary operators needed to determine a complete SI, say *N*_*bo*_, as 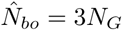, where *N*_*G*_ is the number of genes in the genome of an specie of interest. For example, to determine the complete interactome of *E. coli* we will need a minimum of 3 × 4, 685 = 14, 055 binary operators, while for yeast this figure is 3 × 6, 294 = 18, 882, etc. This naive and rough estimator is likely to be highly biassed, giving smaller number of binary operators than the ones really needed to determine complete SIs; the number of binary operators is more likely to follow an exponential growth as function of the number of genes than a multiplicative one, as assumed above. In [70] the authors presented and demonstrated a general and robust statistical method to estimate the size of interactomes, applying it to protein–protein interactomes, but mentioning that their method can be extended to directed network data, such as gene-regulation networks. The estimation of the sizes of complete SIs using the method presented in [70] will be possible as soon as we have samples of reasonable size of specific SIs and its associated networks which fulfill the sampling requirements asked in that publication.

Finally, in order to apply our algorithmic method to determine and better understand the function of essential cell components, there is a need to merge the broadly disperse interactome data into an integrated SI in which the focus will be the synthesis of cell components. For example, enzymes and metabolic pathways databases, as the one in [95], do not include information about the synthesis of the enzymes from their genetic components, while gene regulatory networks [96] do not include other information, and so forth. Efforts to integrate currently unconnected interactomes in a synthetic framework, as done for example between genomic variant information with structural protein–protein interactomes in [97], or mapping protein–metabolite interactomes as in [98], are the first steps into integrating disperse data. In our opinion, the enormous wealth of disperse interactome knowledge currently existent needs a serious curation effort to obtain integrated SIs, and thus gain further insights about the components essential for life.

## Methods

In this section we present technical concepts that need some definitions and a more precise treatment to be fully explained. However, for brevity we do not present complete formal proofs of our statements.

### Well formed SIs

Synthesis Interactomes (SIs) are structures which contain information about the binary fusion of elements that result into a different element. Table 1 represents a well formed SI of k rows, in which each row is a binary operator of the form ‘*< s*_*ia*_, *s*_*ib*_ *>*⇒ *s*_*i*_’ in which column ‘Name’ contains values *s*_*i*_; *i* = 1, 2, *… k* and column ‘Binary operator’ contains the operator ‘*< s*_*ia*_, *s*_*ib*_ *>*‘. Here we reserve sub-indexed variables, ‘*s*_*i*_, *s*_*ia*_, *s*_*ib*_’ to denote elements of the *i th* row of an SI, while symbols *a, b,* are used for ‘realized’ values of those variables on unspecified rows of an SI. First we will establish that binary operators are commutative, i.e., changing the order of the operands does not change the result, say, if *< a, b >* ⇒ *c* then < *b, a* > ⇒ *c*, thus we have that < *a, b* >=< *b, a* > ⇒ *c*, etc.

The legend of Table 1, gives the conditions for a well formed SI, say i) All represented elements, say *s*_*i*_, *s*_*ia*_, *s*_*ib*_; *i* = 1, 2, *… k*, must be elements of **S**, ii) All names of elements (in column ‘Name’), say *s*_1_, *s*_2_, *…, s*_*k*_, must designate different elements, i.e., *s*_*i*_ ≠ *s*_*j*_ for all pairs *i* ≠ *j* and iii) All *k* binary operators (in column ‘Binary operator’) must be different.

Note that by (i) and (ii) we have that *k* ≤ |**S|** ≤ 3*k*, i.e., the number of elements of **S**, |**S|**, must be of at least *k* and at most 3*k*. This implies that elements that exist as operands in binary operators can also be present in the column ‘Name’, that defines the set of internal elements, **S_i_**, that by (ii) has exactly *k* elements, say |**S_i_**| = *k*. In other words, elements can be repeated within a well formed SI. Condition (iii) implies that if there is a row *r* with value < *a, b* > in column ‘Binary operators’ not other row *i* ≠ *r* could have a value < *a, b* > or < *b, a* >. Also it is worth noting that the order of the rows of an SI is irrelevant; any permutation of rows of an SI will give the same SI and also any not null subset of rows of an SI is a well formed SI.

We also define the set of ‘external’ elements, **S_e_** as **S_e_** = **S** – **S_i_**, the set elements of **S** that do not exist in **S_i_**, and given this there is no synthesis information for them in the SI. Note that 0 ≤ |**S_e_**| ≤ 2*k*. In the previous definition we do not segregate the set of genomic elements, **G**, from the set of external elements. For algebraic manipulations the distinction between **G** and other elements of **S_e_** is only semantic–even if with broad biological relevance, but it has no theoretical consequences for the algorithms used to find essential elements.

### Substitution in binary operators and expanded formulae

An SI defines a finite set of binary operators,

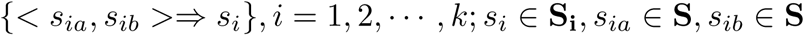

Binary operators can be considered as ‘condensed’ formulae for the synthesis of an element *s*_*i*_. Now we will describe the substitution operation on binary operators that will result into one or more ‘expanded’ formula for the corresponding element. Below we present some relevant definitions.

#### D1 - Substitution in a binary operator

Let < *a, b* > ⇒ *c* be a binary operator into an SI. There are four possibilities for this binary operator, say 1) - *a* ∉ **S_i_** and *b* ∉**S_i_**; 2) - *a* ∉**S_i_** and *b* ∈ **S_i_**; 3) - *a* ∈ **S_i_** and *b* ∉ **S_i_** and 4) - *a* ∈ **S_i_** and *b* ∈ **S_i_**. A substitution in a binary operator is defined as the replacement in the binary operator of the operands by their corresponding binary operators, when they exist. The string resulting from this operation will be called the ‘expanded formulae of level 1’, and for any *x* ∈ **S_i_** will be denoted by ε_1_(*x*). We also define the expanded formula of order 0, say ε_0_(*x*), as the binary operator for *x*.

#### D2 - Substitution in an expanded formula

Let *ε*_*r*_(*x*) be an expanded formula for *x*, and 𝒪_*r*_(*ε*_*r*_(*x*))) denote set of operands in this formulae, i.e., 𝒪_*r*_(*ε*_*r*_(*x*))) is the set of all symbols that represent elements of **S** within the formula *ε*_*r*_(*x*). The expanded formula of order *r* + 1 for *x*, say, *ε*_*r*+1_(*x*), is defined as the result of substituting all elements of **S_i_** ∈ 𝒪_*r*_(*ε*_*r*_(*x*))) by their corresponding binary operators in *ε*_*r*_(*x*).

#### D3 - The complete set of operands of order *r* for *x*

We define the ‘complete set of operands of order *r* for *x* for an *x* ∈ **S_i_** as

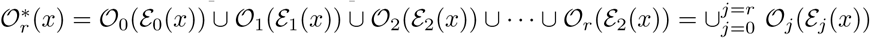

#### D4 - A closed expanded formula

We define a closed expanded formula for an *x* ∈ **S_i_**, say *ε\**(*x*) = *ε*_*r*_(*x*), as the expanded formula for *x* such that *ε*_*r*_(*x*) *≡ ε*_*r*+1_(*x*) if there is a value of *r*; *r* = 1, 2, *…* such that the condition *ε*_*r*_(*x*) *≡ ε*_*r*+1_(*x*) is fulfilled.

The definitions above imply that we can proceed in consecutive steps, say *r* = 0, 1, 2, *…*, to obtain expanded formulae from the synthesis information present in the SI. **D1** defines *ε*_0_(*x*) as the binary operator for *x* and gives the method to obtain *ε*_1_(*x*). It is clear that if both operands in *ε*_0_(*x*) are external structures in **S_e_** then *ε*_1_(*x*) *=ε*_0_(*x*), simply because there is no element to be substituted and the ‘expanded’ formula for *x* will be in that case identical to the binary operator, *ε*_0_(*x*). **D2** explains the procedure to obtain *ε*_*r*+1_(*x*) from the formula obtained in the previous step, *ε*_*r*+1_(*x*), completing the method to obtain the sequence

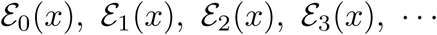

which is a nested process of substitution, which expands all information existent into the SI for the synthesis of *x*. To be able to define *ε*_*r*+1_(*x*) as function of *ε*_*r*_(*x*), **D2** also defines the set of operands present into a formula, say, 𝒪_*r*_(*ε*_*r*_(*x*))). Clearly, the only elements of 𝒪_*r*_(*ε*_*r*_(*x*))) that could be substituted by their binary operators, are internal elements in **S_i_**. This implies that if 𝒪_*r*_ *∩* **S**_**i**_ = ϕ then ε_*r*+1_(*x*) ≡ε_*r*_(*x*), i.e., not change will be produced in the expanding formula, because no substitution was performed. That in turn means that the formula _*r*_(*x*) for *x* is a ‘closed expanded formula’, as defined in **D4**. This can be summarized as a first theorem,

#### T1. Existence of a closed expanded formula for *x*

A closed expanded formula for *x* ∈ **S_i_** exist if and only if for a given value of *r* the condition 𝒪_*r*_(*ε*_*r*_(*x*))) ∩ **S_i_** = *ϕ* is fulfilled. In such case *E*_*r*_(*x*) is a closed expanded formula for *x*, that will be denoted as *ε\**(*x*).

The proof of this theorem is obtained by showing the necessity and sufficiency of the condition 𝒪_*r*_(*ε*_*r*_(*x*))) ∩ **S_i_** = *ϕ*.

A first consequence of **T1** is that closed formulae are formed exclusively by external elements. This is obvious because if 𝒪_*r*_(*ε*_*r*_(*x*))) ∩ **S_i_** = *ϕ* is true, then 𝒪_*r*_(*ε*_*r*_(*x*))) ∩ **S_e_** = 𝒪_*r*_(*ε*_*r*_(*x*))) given that all elements of 𝒪_*r*_(*ε*_*r*_(*x*))) are elements of **S** and **S** = **S_i_** ∪ **S_e_**; **S_i_** ∩ **S_e_** = *ϕ*.

Even when **T1** gives the condition for the existence of a ε*(*x*) for *x*, it does not in general guarantee the existence of such closed formula for *x*, thus it is possible that the sequence *ε*_0_(*x*), *ε*_1_(*x*), *ε*_2_(*x*), *…* will never provide such formula, if the condition in **T1** is not fulfilled. In fact, the negation of the condition 𝒪_*r*_(*ε*_*r*_(*x*))) *=* ϕ, say, that there is not a value of *r* = 0, 1, … for which this condition is fulfilled, implies the possibility of elements *x* for which there are only ‘open’ expanded formulae. To analyze those cases, let examine the definition of the complete set of operands of order *r* for *x*, denoted as 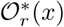 and defined in **D3** as 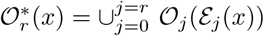.

First, we can say that for any *x* ∈ **S_i_** we have that 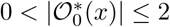, i.e., the number of elements of this set will be the number of distinct operands in the binary operator corresponding to *x*, and this can only be 1 if both operands are the same, or 2 if they are different, given that 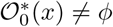. Second, it is clear that 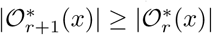, because re-writing the definition in **D3** we have

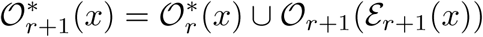

i.e., the number of elements in the set 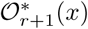 cannot decrease, and will stay the same, that is 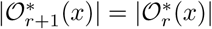 if and only if 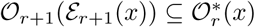, i.e., if no more elements are added to the set 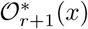 by the union with 𝒪_*r*+1_(ε _*r*+1_(*x*)).

It appears to be clear that the set 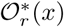 cannot grow indefinitely, and in fact its maximum size, 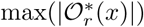, cannot be larger that the number of elements named in the corresponding SI, say

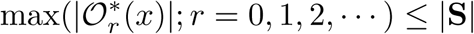

From this we can postulate the following theorem

#### T2. Convergence of the complete set of operands

For every element *x* ∈ **S_i_** there is a value *u* ∈ {0, 1, 2, …, *z* – 1, *z*} such that

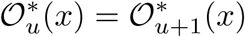

A *reductio ad absurdum* proof of **T2** results directly from the fact that

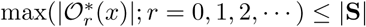, because given that 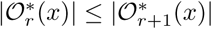, there must exist the number *u*–as postulated in **T2**, for which 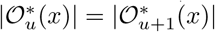 and in that case 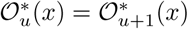, because for all *r* we have that 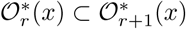. Assuming that there is not a value of *u* that fulfills **T2** leads to a contradiction.

Let’s briefly give some details. Assume that for *x* ∈ **S_i_** there exist a closed expanded formula, *ε\**(*x*), and denote by *u* the smallest number that fulfills 𝒪_*u*_(*ε*_*u*_(*x*)) ∩ **S_i_** = *ϕ*. A consequence of the existence of a closed expanded formula for *x* is that *ε*_*u*+1_(*x*) *=ε*_*u*_(*x*), and by induction also *ε*_*u*+*j*_(*x*) ≡ *ε*_*u*_(*x*); *j* = 1, 2, … In other words, after the point *u* the expanded formula for *x* does not change, and this in turn means that the set 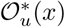 will not have any additional elements from operands in further expanded formulae, *εu*+1(*x*), *ε*_*u*+2_(*x*), … demonstrating that, for elements with a closed expanded formula, *u* is the point mentioned in **T2**.

Now, let’s take the case of *x* ∈ **S_i_** for which there is not a closed expanded formula. In that case the expanded formula for *x* will be always increasing in the number of terms as function of the number of substitution steps. To be specific, denote as *T* (*ε*_*r*_(*x*)) the function that gives the total number of symbols included into the expanded formula *ε*_*r*_(*x*). Given that there is not a closed expanded formula for *x* it follows that *T* (*ε*_*r*_(*x*)) < *T* (*ε*_*r*+*j*_(*x*)) for *j* = 1, 2, *…*; in words, we will have a never ending increase in the number of symbols forming *ε*_*r*_(*x*) as the number of substitution steps increases. However, while *T* (*ε*_*r*_(*x*)) is not bounded, the number of elements in 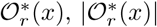 is in fact limited; we have seen that the absolute maximum for 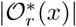 is *|***S***|*. Thus, to find the value of *u* in **T2** for cases where *x* does not have a closed expanded formula we need to algorithmically find the smallest value of *r* that fulfills the condition 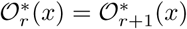, and this value must exist because 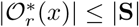.

Assume that we have found the value of *u* for the case of *x* ∈ **S_i_** with no closed formula; i.e., in a particular case we corroborate that 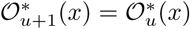–as postulated by **T2**. We only need to see that for any value *u* + *j*; *j* = 2, 3, *…* the equality 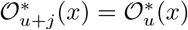 holds for all values of *j* = 2, 3, *…*. But that is clear because 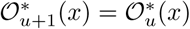 means that at step *u* there was no new internal elements of *S*_*i*_ to be substituted into *ε*_*u*+1_(*x*), i.e., all elements of *S*_*i*_ that could be operands in any *εk* (*x*); *k < u* had been already found in a previous step and thus they are already into the set 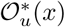; that is why there is not change from 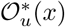 to 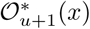. Thus, we can simplify our notation and denote the complete set of operands for *x* simply as 𝒪*(*x*), understanding that this is the larger and stable set which will not depend on *u*.

We have seen that in general, for every *x* ∈ **S_i_** we can find a number of nested substitutions, *u*, for which 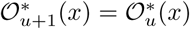, independently if the element *x* has a closed (*ε\**(*x*)) or open formula. Now we can define the central property of ‘recursiveness’ of an element *x*, from which we infer biological essentiality.

#### D5. Recursiveness of an structure *x*

An element *x* ∈ **S_i_** is said to be recursive if an only if 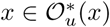, where *u* is the smallest integer for which 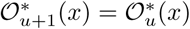.

In the main text we have discussed why if an element is recursive then it is also essential for the cell, leading to our first rule of essentiality, **ER1**. The second rule of essentiality, **ER2**, also discussed at the main text, says that all operands found in 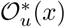 for a recursive element *x*, are also essential.

To exemplify the definitions given above and appreciate their consequences, Table 4 presents a simple SI.

**Table 4.** A simple SI (including extra column ‘Row’ for reference).

| Row | Name | Binary operator |
| --- | --- | --- |
| 1 | $a$ | $\langle b, c \rangle$ |
| 2 | $b$ | $\langle a, d \rangle$ |
| 3 | $f$ | $\langle e, h \rangle$ |
| 4 | $g$ | $\langle f, f \rangle$ |
| 5 | $i$ | $\langle a, b \rangle$ |

For this SI we have: **S** = {*a, b, c, d, e, f, g, h, i*}; **S_i_** = {*a, b, f, g, i*}; **S_e_** = {*c, d, e, h*}.

For the SI presented in Table 4 we can easily see that for the row 3, which defines the synthesis of *f* by the binary operator < *e, h* > (< *e, h > ⇒ f*) the substitution in the binary operator has no effect, given that both operands, *e* and *h*, are external structures (in **S_e_**) and thus we have that *ε*_*r*_(*f*) =< *e, h* > for *r* = 0, 1, 2, and also, for any value of *r* we have that 𝒪_*r*_(*ε* _*r*_(*f*)) = {*e, h*} and 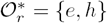, thus there exist a closed expanded formula for *f* which in this case is simply given by < *e, h* >. This is illustrated in Table 5.

**Table 5.** Expressions for element *f* of the SI in Table 4 (row 3 in that table).

| $r$ | $\mathcal{E}_r(i)$ | $\mathcal{O}_r(\mathcal{E}_r(i))$ | $\mathcal{O}_r^*(i)$ |
| --- | --- | --- | --- |
| 0 | $\langle e, h \rangle$ | $\{e, h\}$ | $\{e, h\}$ |
| 1 | $\langle e, h \rangle$ | $\{e, h\}$ | $\{e, h\}$ |
| 2 | $\langle e, h \rangle$ | $\{e, h\}$ | $\{e, h\}$ |
| … | $\langle e, h \rangle$ | $\{e, h\}$ | $\{e, h\}$ |

A more interesting case, where we can observe the consequences of the definitions given above happens with the element *i* given in the row 5 of Table 4. Table 6 presents the values of *ε*_*r*_(*i*), 𝒪_*r*_(*ε*_*r*_(*i*)) and 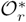 for different values of *r*.

**Table 6.**
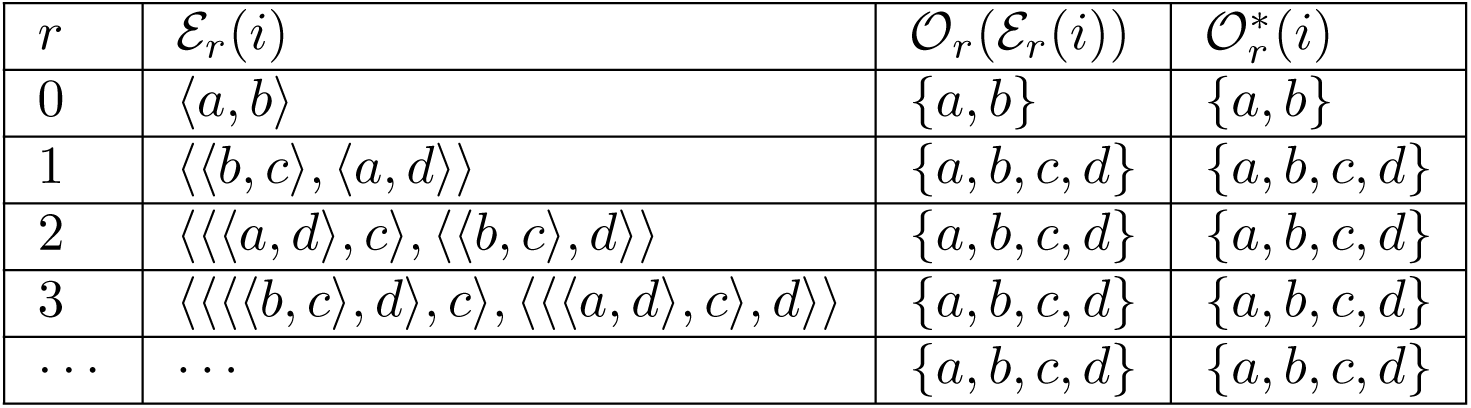
Expressions for element *i* of the SI in Table 4 (row 5 in that table).

In Table 6 we can see how the expanded formula for *i, ε*_*r*_(*i*), continues expanding as *r* increases. In fact, when *r* = 10 the number of symbols present in ε_10_(*i*) is of 93 (data not shown), etc. For this case an algorithm to continue substituting into a formula ‘until it stops growing’ will fall into an infinite loop. In contrast, the set of operands for the formula, 𝒪*ε*_*r*_(*ε*_*r*_(*i*))–the set of operands in the formula *ε*_*r*_(*i*), as well as 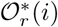–the set of operands that have appeared in any of the steps (including the current one), are stabilized as *a, b, c, d* after the first substitution, i.e., for *r* = 2, 3,. From this we conclude that there is not a closed expanded formula for *i*, i.e., it is not posible to find a value of *r* for which *ε*_*r*_(*i*) ≡ *ε*_*r*+1_(*i*) is fulfilled.

Now let’s examine the expressions for the expansion of the formula of *a* (row 1 of Table 4), presented in Table 7.

**Table 7.** Expressions for element *a* of the SI in Table 4 (row 1 in that table).

| $r$ | $\mathcal{E}_r(a)$ | $\mathcal{O}_r(\mathcal{E}_r(a))$ | $\mathcal{O}_r^*(a)$ |
| --- | --- | --- | --- |
| 0 | $\langle b, c \rangle$ | $\{b, c\}$ | $\{b, c\}$ |
| 1 | $\langle \langle a, d \rangle, c \rangle$ | $\{a, c, d\}$ | $\{a, b, c, d\}$ |
| 2 | $\langle \langle \langle b, c \rangle, d \rangle, c \rangle$ | $\{b, c, d\}$ | $\{a, b, c, d\}$ |
| 3 | $\langle \langle \langle \langle a, d \rangle, c \rangle, d \rangle, c \rangle$ | $\{a, c, d\}$ | $\{a, b, c, d\}$ |
| 4 | $\langle \langle \langle \langle \langle b, c \rangle, d \rangle, c \rangle, d \rangle, c \rangle$ | $\{b, c, d\}$ | $\{a, b, c, d\}$ |
| … | … | … | $\{a, b, c, d\}$ |

From the first 4 rows of Table 7 we can infer that there is not a closed expanded formula for the element *a* of the SI presented in Table 4; the process of substitution can continue without ever arriving at a value of *r* such that *ε*_*r*_(*a*) ≡ *ε*_*r*+1_(*a*) is fulfilled. On the other hand we can also see that the sets of operands that appear in the expanded formula of order *r* [𝒪_*r*_(*ε*_*r*_(*i*)) in the third column of Table 7] do not stabilize, varying from {*b, c*} for *r* = 0, {*a, c, d*} for *r* = 1, {*b, c, d*} for *r* = 2 and then alternating between these two values at consecutive rows. In contrast, the set of operands that have appeared in any of the steps (including the current one), 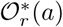 is stable as {*a, b, c, d*} after the first substitution (at *r* = 1).

The fact that the set 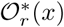, obtained as examples for the elements *f, i* and *a* of the SI presented in Table 4 ‘stabilizes’ after a number of iterations indicates that we have substituted all internal elements in **S_i_** at the corresponding formula.

### An algorithm to find essential elements

Here we summarize the algorithm to find all recursive structures within the ones defined in a well formed SI. The basic idea is to keep performing nested additions of members to the sets of operands for each *s*_*i*_; *i* = 1, 2, *…, k* until obtaining all sets of complete operands, 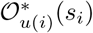; *i* = 1, 2, *…, k*–that is, until obtaining all the stable sets of operands for each *s*_*i*_ as defined in **T2** (note that *a priory* the values of *u* could be different for each *i*, and therefore are denoted as ‘*u*(*i*)’ in the previous expression).

Having the collection 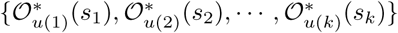, we can examine for which cases we have that 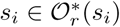, i.e., we can determine which elements have a recursive set of operands (see **D6**), and therefore will fulfill the first essentiality rule **ER1**.

The key aspect for the implementation of the algorithm is to perform additions only of elements which are not recursive, otherwise the procedure could fall into an infinite loop, trying a never ending chain of nested additions to some sets. A problem is that *a priory* we do not know which elements have a recursive formula. A solution (found by M. H. R-V) is to mark as ‘frozen’ those elements which are found to have a recursive formula as soon as they are detected, and from then on avoid the substitution of the set for such ‘frozen’ elements into subsequent steps. The algorithm ends when all sets of operands are stable or ‘complete’ as demanded by **T2**.

In practice various rounds of addition could be needed for the algorithm to be completed; note that here the word ‘addition’ means to include an element into a set, but if such element is already in the set, it will not increase the size of such set. This procedure begins by assuming that there are not recursive elements and thus a logical vector ‘`frozen`’ of *k* elements is defined as ‘`FALSE`’ for *i* = 1, 2, *k*. Also a list of *k* ‘`current`’ sets is initialized by setting ‘`current[i]`’ equal to the set of the corresponding operands, i.e., if *i* = 1 and the first row of the SI contains `“a”` and `“b”` in columns *o*1 and *o*2 respectively, then ‘current[1] = (“a”, “b”)’, etc.

After the initialization of the ‘`current`’ vector it is checked to find if any of its elements must be frozen. This is done by testing if the name of the internal structure, the column ‘Name’ of the SI in a vector of *k* elements ‘`name`’, is a member of the corresponding set of operands, i.e., if ‘`name[i] 〈 current[i]`’. For all elements that fulfill such condition, i.e., recursive elements, the corresponding value of ‘`frozen`’ is set to ‘`TRUE`’. This process of ‘frozen update’ will be repeated after each round of additions to the sets.

The next step is to perform the creation of new sets after adding elements. For this a list of *k* new sets is defined as ‘`new`’. To find the elements of each ‘`new`’ set, say `new[i]; i = 1, 2, … k`, each one of the elements of the corresponding ‘`current[i]`’ vector (current[i][1], current[i][2], …) are analyzed and the procedure in list **(i)** is applied.

#### (i)- Obtain new sets from current ones

1. If `current[i][j]` ∉**S_i_** then `current[i][j]` is included into `new[i]`. Otherwise,
2. If `current[i][j]` is ‘frozen’ (marked as recursive) then `current[i][j]` is included into `new[i]` without performing a substitution of its operands. Otherwise,
3. At this point we know that `current[i][j]` is an internal structure (∈ **S_i_**) which is not frozen, thus an addition of members to ‘`new[i]`’ must be performed. We look which element of ‘`name`’ is equal to `current[i][j]`. Say that ‘`name[k] = current[i][j]`’, then we include all elements of ‘`current[k]`’ into ‘`new[i]`’.

At this point we can test if all sets in the lists ‘`current`’ and ‘`new`’ are equal. If that is the case it means that we have found all sets of complete operands defined in **T2**. Otherwise we must iterate the procedure shown in list **(i)** as many times as necessary to obtain all sets of complete operands. The procedure in in list **(ii)** below must be performed until “all sets in the lists ‘`current`’ and ‘`new`’ are equal”.

#### (ii)- Iterate until all sets new and current are identical

1. Set ‘`current = new`’.
2. Update the value of the ‘`frozen`’ vector (see if new recursive structures are detected and froze them).
3. Obtain a new value for the list ‘`new`’ (procedure in list **(i)**).

By running the procedure described above until convergence (lists **(i)** and **(ii)**), and reviewing which elements are recursive, we could obtain a list of sets 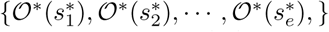, where each of the elements 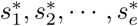 is essential, given that 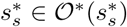 *s* = 1, 2, *…, r*.

The second rule of essentiality, **ER2**, enunciated before in the main text, states that all operands which intervene in the synthesis of an essential structure are also essential. Thus, to obtain the complete set of essential structures for the SI, say, **E**, we must perform the union of each one of the sets 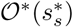; *s* = 1, 2, *…, r*, that is

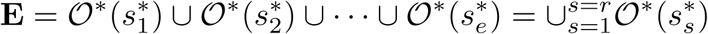

Naturally it could happen that **E** = *ϕ*, i.e., it was not possible to determine any essential structure for the SI or, on the other extreme, **E** = **S**, i.e., all structures named within the SI are judged to be essential. As discussed in the main text, ‘essentiality’ is only judged within the framework of the information contained in the corresponding SI.

The algorithm presented here is implemented in the R environment [72] within our package ‘InterPlay’ (included as ‘S1 Binary’) and amply exemplified in ‘S1 Text’. Also, plotting of SIs as networks is exemplified in that appendix (S1 Text).

## Supporting information

**S1 Text. Additional text and computational examples.** Additional details and discussion of the C2E algorithm applied to the cases of the RNA polymerase and streptomycin SIs. Also demonstrates the functions and data of our R [72] package ‘InterPlay’ (included as ‘S1 Binary’) to work with SIs and determine essential structures. Algorithms are exemplified and explained in detail and plotting of interactome networks is illustrated with the use of the ‘igraph’ [73] R package (see also S2 Text).

**S2 Text. Supplementary functions.** R functions to plot SIs as networks using our package ‘InterPlay’ (included as ‘S1 Binary’) as well as the ‘igraph’ [73] R package.

**S3 Text. InterPlay manual.** Manual for our R package ‘InterPlay’ (included as ‘S1 Binary’).

**S1 Binary. InterPlay R package.** Binary file with our R package ‘InterPlay’. The manual for this package is presented as ‘S3 Text’. To install this R [72] package see ‘S1 Text’ or the corresponding R documentation.

## Acknowledgments

We are grateful to Gerardo R. Argüello-Astorga, June Simpson and Therese A. Markow for useful discussion and suggestions as well as to three anonymous referees for valuable suggestions and criticisms.

